# Conservation of the ancestral function of GTSF1 in transposon silencing in the unicellular eukaryote *Paramecium tetraurelia*

**DOI:** 10.1101/2023.10.06.561219

**Authors:** Chundi Wang, Liping Lv, Therese Solberg, Zhiwei Wen, Haoyue Zhang, Feng Gao

## Abstract

The PIWI-interacting RNA (piRNA) pathway is crucial for transposon repression and the maintenance of genomic integrity. Gametocyte specific factor 1 (GTSF1), an indispensable auxiliary factor of PIWI, was recently shown to potentiate the catalytic activity of PIWI in many metazoans. Whether the requirement of GTSF1 extends to PIWI proteins beyond metazoans is unknown. In this study, we identified a homolog of GTSF1 in the unicellular eukaryote *Paramecium tetraurelia* (PtGTSF1) and found that its role as a PIWI-cofactor is conserved. PtGTSF1 interacts with PIWI (Ptiwi09) and Polycomb Repressive Complex 2 (PRC2) and is essential for PIWI-dependent DNA elimination of transposons during sexual development. PtGTSF1 is crucial for the degradation of PIWI-bound small RNAs recognizing the organism’s own genomic sequences. Without PtGTSF1, self-matching small RNAs are not degraded and results in an accumulation of H3K9me3 and H3K27me3, which disturbs transposon recognition and slows down their elimination. Our results demonstrate that the PIWI-GTSF1 interaction also exists in unicellular eukaryotes with the ancestral function of transposon silencing.

## Introduction

Transposable elements (TEs) are found in the genomes of nearly all organisms, and can make up a significant portion of the genome (Lawson *et al*, 2023). They contribute to genetic diversity and evolution by causing mutations, driving gene duplication, and creating new genes (Modzelewski *et al*, 2022). However, their uncontrolled activity can lead to genomic instability. Therefore, most organisms have evolved mechanisms to silence or limit the activity of TEs. The PIWI-interacting RNA (piRNA) pathway is essential for the repression of TEs in animal germlines as well as in some somatic tissues of various non-mammalian species (Jehn *et al*, 2018; Lewis *et al*, 2018; Wang *et al*, 2023). piRNAs are transcribed from genomic loci known as piRNA clusters, which are further processed into primary piRNAs with the ability to guide its PIWI-partner to TE transcripts (Aravin *et al*, 2007; Brennecke *et al*, 2007). Cleavage by a PIWI protein initiates the production of secondary piRNAs that can further participate in TE repression (Mohn *et al*, 2014; Wang *et al*, 2023). Transcriptional TE silencing is achieved by nuclear PIWI proteins (*e.g.,* Piwi in flies and MIWI2 in mice), which are guided by piRNAs to nascent TE transcripts and generate heterochromatin via DNA or histone methylation (*e.g.,* trimethylation of lysine 9 on histone H3, H3K9me3). Post-transcriptional TE silencing is mediated by cytoplasmic PIWI proteins (*e.g.,* Aub and Ago3 in flies, MILI and MIWI in mice), which can directly slice target transcripts complementary to their bound piRNA (as reviewed in Ozata *et al*, 2019).

Ciliates, one of the most diverse groups of unicellular eukaryotes that emerged approximately one billion years ago, employ a more extreme approach of controlling TEs than most organisms, and do not rely on constant transcriptional or post-transcriptional suppression of such elements. They separate germline and somatic functions into distinct nuclei in the same cytoplasm, and only allow TEs and TE-derived sequences known as internally eliminated sequences (IESs) to exist in the transcriptionally inactive germline micronuclear (MIC) genome. During sexual processes, they are eliminated from the active somatic macronuclear (MAC) genome through a piRNA-like DNA elimination pathway (as reviewed in Gao *et al*, 2023; Rzeszutek *et al*, 2020). In the ciliate *Paramecium tetraurelia* (*P. tetraurelia*), about one third of germline-limited sequences are eliminated during sexual processes, including TEs and ∼45,000 IESs (Arnaiz et al., 2012).

The piRNA-like pathway in *P. tetraurelia* involves a small RNA-mediated comparison of the MIC and MAC genomes, known as the “scanning model” (Mochizuki *et al*, 2002). During this process, small RNAs generated from the germline MIC genome “scan” the somatic MAC genome to remove self-matching sequences. This enriches for small RNAs containing MIC-specific sequences (including TEs and IESs), which are used to guide their elimination in the developing new MACs. When the sexual process is initiated, the whole MIC genome is bidirectionally transcribed by RNA polymerase II to produce double-stranded long noncoding RNAs (dsRNAs), which are cleaved into 25 bp small noncoding RNAs (scnRNAs) by Dicer-like proteins (Lepere *et al*, 2009; Sandoval *et al*, 2014). Double-stranded scnRNAs are transported to the cytoplasm where they are loaded onto PIWI proteins (Ptiwi01/Ptiwi09) and the “passenger” strand is removed (Bouhouche *et al*, 2011). The PIWI-scnRNA complexes are transported to the old MAC where they “scan” nascent transcripts by homologous pairing and those that find complementary sequences are degraded by an unknown mechanism. The remaining scnRNAs containing MIC-specific sequences are transported to the developing new MACs where they pair with nascent transcripts and recruit Polycomb repressive complex 2 (PRC2) to mediate the deposition of H3K9me3 and H3K27me3 (Miró-Pina *et al*, 2022; Wang *et al*, 2022). These modifications help recruit the domesticated PiggyBac transposase PGM that excises the MIC-specific DNA (Baudry *et al*, 2009; Bischerour *et al*, 2018). After excision, the excised DNA initiates the production of a secondary class of small RNAs known as iesRNAs, which serve as a positive feedback loop to ensure complete elimination of all MIC-specific DNA (Allen *et al*, 2017; Furrer *et al*, 2017; Sandoval *et al*, 2014). Finally, the remaining DNA segments (called macronuclear destined sequences, MDSs) are ligated by non-homologous end joining (NHEJ) to form the new somatic MAC genome (Kapusta *et al*, 2011; Marmignon *et al*, 2014).

Although this model was proposed 20 years ago, and an increasing number of factors involved in this process have been identified, many questions remain unanswered. One of these questions is how self-matching MDS-scnRNAs are selected and degraded, to prevent their entry into the new MACs where they could guide PRC2 and lead to errant DNA elimination. In this study, we approached this question by identifying and characterizing a PIWI- and PRC2-interacting protein essential for this process. This protein is a homolog of Gametocyte-specific factor 1 (GTSF1), an evolutionarily conserved small zinc finger protein, which has been reported in many animals (Almeida *et al*, 2018; Chen *et al*, 2020; Dönertas *et al*, 2013; Ohtani *et al*, 2013; Yoshimura *et al*, 2009). GTSF1 was first identified in unfertilized eggs, ovaries, and testes in mouse, and later revealed to be an essential component of the piRNA pathway in various organisms, including mouse, *Drosophila* and silkworm (Chen *et al*, 2020; Dönertas *et al*, 2013; Muerdter *et al*, 2013; Ohtani *et al*, 2013; Yoshimura *et al*, 2007; Yoshimura *et al*, 2009; Yoshimura *et al*, 2018). GTSF1 directly interacts with both cytoplasmic and nuclear PIWI proteins and is essential for transcriptional and post-transcriptional regulation of transposable elements. A recent study revealed that GTSF1 enhances the intrinsically weak RNA cleavage activities of catalytically-active PIWI proteins (Arif *et al*, 2022).

In this study, we identified a homolog of GTSF1 in the unicellular eukaryote *P. tetraurelia* (PtGTSF1) and found the relationship between GTSF1 and PIWI to be conserved. PtGTSF1 plays a crucial role in the repression of TEs by facilitating the degradation of scnRNAs recognizing the organism’s own genomic sequences, a mechanism which distinguishes self- from non-self-DNA. Our results demonstrate that PtGTSF1 interacts with both PIWI and the PRC2 complex, and regulates the scnRNA pool that guides the PRC2 complex to mark sequences for elimination. The identification of GTSF1 in protists indicates that GTSF1 was present in the last common ancestor of all animals with the conserved function of RNA-mediated transposon silencing.

## Results

### PtGTSF1 interacts with Ptiwi09 and PRC2

In a previous study, we performed immunoprecipitation and mass spectrometry to identify the PRC2 complex and its interaction partners (Wang *et al*, 2022). In addition to identifying the subunits of the complex, we discovered a homolog of gametocyte-specific factor 1 in *P. tetraurelia* (PtGTSF1) (Fig EV1A-D). We also re-analyzed mass spectrometry data from a related study in which the interaction partners of the scnRNA-binding PIWI protein Ptiwi09 was reported and found PtGTSF1 to also co-precipitate with Ptiwi09 (Fig EV1E) (Miró-Pina *et al*, 2022). Since GTSF1 is an essential component of the piRNA pathway in animals and is well known to interact with PIWI proteins, we decided to further characterize this protein (Almeida *et al*, 2018; Arif *et al*, 2022; Dönertas *et al*, 2013; Ohtani *et al*, 2013; Yoshimura *et al*, 2018).

PtGTSF1 (PTET.51.1.P0490019 in ParameciumDB) is predicted to have two CHHC-type zinc fingers, which is the hallmark of GTSF1 proteins (Fig 1B and Appendix Fig S1) (Arnaiz *et al*, 2020). Although mammals have three GTSF1 paralogs (GTSF1, GTSF1-like, and GTSF2) and *Drosophila* has four (Asterix/DmGTSF1/CG3893, CG14036, CG32625, and CG34283), we did not identify any PtGTSF1 paralog in *P. tetraurelia* by blastp (Muerdter *et al*, 2013; Takemoto *et al*, 2016). PtGTSF1 clusters with *Drosophila melanogaster* and *Caenorhabditis elegans* GTSF1 rather than their paralogs, indicating that PTET.51.1.P0490019 is GTSF1 in *P. tetraurelia* (Fig 1C). To further confirm the interactions identified in the previously published mass spectrometry data, we expressed a HA-tagged PtGTSF1 in *P. tetraurelia* and performed immunoprecipitation using an anti-HA antibody to identify interaction partners of PtGTSF1 (Fig EV1G). Both Ptiwi09 and PRC2 subunits were co-immunoprecipitated with PtGTSF1, confirming the interactions (Fig EV1F). Considering the previously established interaction between Ptiwi09 and PRC2 (Miró-Pina *et al*, 2022; Wang *et al*, 2022), we conclude that PtGTSF1, PRC2 and Ptiwi09 interact in *P. tetraurelia* (Fig EV1H).

**Figure 1.**
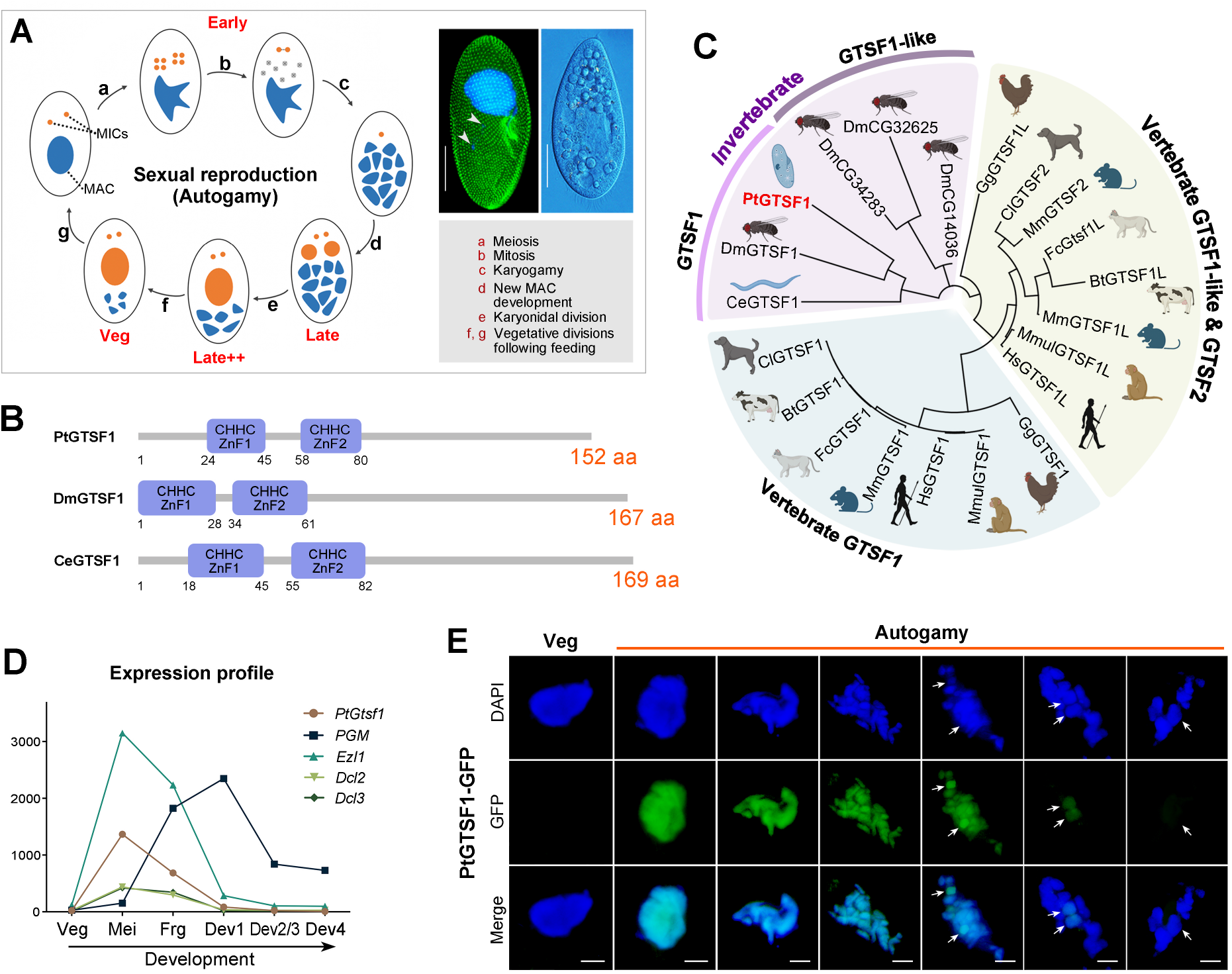
PtGTSF1 is exclusively expressed during sexual development in Paramecium tetraurelia. A Photographs and sexual reproduction (autogamy) in *P. tetraurelia*. (a) Two micronuclei (MICs) undergo meiosis to produce eight haploid nuclei. (b) One of eight haploid nuclei divides mitotically and the rest degrade. (c) Two haploid nuclei fuse to form the zygotic nucleus. (d) The zygotic nucleus divides twice before two products become the new MICs and the other two develop into the new Macronuclei (MACs). (e) The cell divides once, distributing the nuclei into two cells. (f, g) Refeeding after autogamy recovers the cell to the vegetative stage. The timepoints collected in this study are labeled in red (Early, Late, Late++ and Veg). Arrowheads point to micronuclei. Scale bar: 30μm. B Protein domains of GTSF1 in *Paramecium tetraurelia* (Pt), *Drosophila melanogaster* (Dm), and *Caenorhabditis elegans* (Ce). C Neighbor-Joining tree of GTSF1 and its paralogs in *P. tetraurelia* and metazoans. D RNA-seq expression profiles of *PtGtsf1*, *PGM*, *Ezl1*, *Dcl2* and *Dcl3*. Veg: vegetative growth; Mei: meiosis; Frg: fragmented old MAC; Dev1-4: stages of new MAC development. E Localization of PtGTSF1 during sexual development (autogamy). From left to right, progression of autogamy. Arrows indicate new MACs. Scale bar: 10μm.

### PtGTSF1 is essential for sexual development

The life cycle of *Paramecium* is separated into vegetative and sexual stages. According to publicly available mRNA-seq data, *PtGtsf1* is specifically expressed during sexual development (autogamy), with the highest levels detected at the meiotic stage (Fig 1A and D) (Arnaiz *et al*, 2020). This expression pattern is reminiscent of genes encoding core components of PRC2 and scnRNA-related proteins, such as Ptiwi01 and Ptiwi09, and Dicer-like proteins Dcl2 and Dcl3 (Bouhouche *et al*, 2011; Sandoval *et al*, 2014).

We next assessed the localization of PtGTSF1 by fusing it with a green fluorescent protein (GFP) (Fig 1E). No fluorescence was observed during vegetative growth. Upon induction of autogamy by starvation, PtGTSF1 initially appeared in the intact old MAC and remained there when the MAC threaded and fragmented. In early new MAC development, PtGTSF1 exists both in the old and new MACs simultaneously. As the new MACs continue to develop, the signal gradually diminishes, leaving only faint signals in the new MACs which eventually disappear. Through the entire life cycle, PtGTSF1 was never detected in the MICs.

Both the expression profile and protein localization suggest that PtGTSF1 has a function specifically during sexual development in *P. tetraurelia*. To further investigate its function, we performed knockdown of *PtGtsf1* (*PtGtsf1* KD) and an empty vector (EV) control. During the vegetative stage, *PtGtsf1* KD cells grow similarly to the EV control, confirming that it is dispensable for vegetative growth. To accurately assess its function during the autogamy process, we studied four representative phases (Fig 1A). Early, corresponding to the meiotic stage, in which 30%–50% of cells have a fragmented old MAC. Late, approximately 16 hours after Early with more than 70% of cells having visible new MACs. At this stage, TEs and IESs in the new MACs are being eliminated. Late++, 36 hours after Late when the new MACs are almost fully developed. Veg, 24 hours after feeding Late++ cells, corresponding to the vegetative stage of next life cycle.

*PtGtsf1* KD cells can enter autogamy and there are no noticeable differences compared to the control until the Late++ stage, at which point the cells have an increased number of old MAC fragments compared to the control (median: 11.0 vs. 22.5), and this difference remains significant until the Veg stage (4.0 vs. 11.5) (Fig 2A and B). Unlike the control, *PtGtsf1* KD cells cannot recover to the vegetative stage after autogamy. After re-feeding, *PtGtsf1* KD cells survive the first day; however, their morphology is aberrant (Fig 2C). The posterior end changed from broadly rounded to irregular, and the cells became shorter and wider with a decreased length/wide ratio (Fig 2C and D). From the second day, cell numbers in the *PtGtsf1* KD culture show obvious differences to the control with nearly all of them sick or dead (Fig 2E). Taken together, the completion of sexual development, but not vegetative growth, requires PtGTSF1 in *P. tetraurelia*.

**Figure 2.**
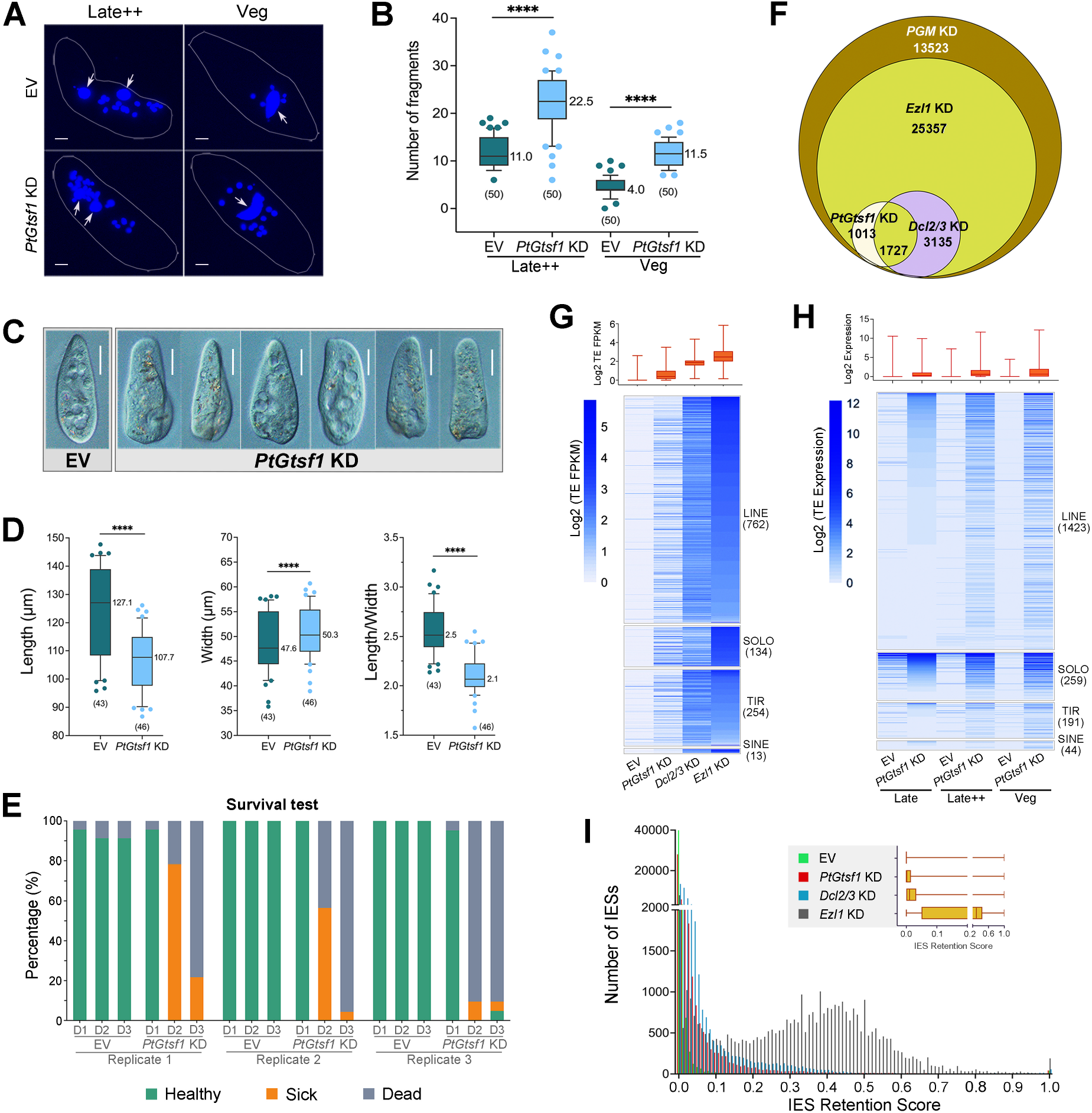
PtGTSF1 is essential for sexual development and the repression of TEs. A DAPI staining to show the old MAC fragments in Late++ and Veg stages of EV (empty vector, negative control) and *PtGtsf1* KD. Arrows indicate new MACs. Scale bar: 10μm. B Numbers of old MAC fragments in EV and *PtGtsf1* KD. Numbers in brackets under the boxes are the numbers of individuals whose fragments were counted. Horizontal line and the number to the right of the line denotes the median. **** p < 0.0001 (unpaired t-test). C Photographs of *P. tetraurelia* on the first day after feeding. Scale bar: 30μm. D The length, width, and length/width ratio of *P. tetraurelia* cells on the first day after feeding. E Survival test of EV and *PtGtsf1* KD cells. Cells were monitored for three consecutive days after feeding, denoted by D1, D2 and D3. Green: healthy; Orange: sick; Gray: dead. F Venn diagram depicting the overlap of IESs (IES retention scores ≥0.1) affected by *PtGtsf1* KD, *Dcl2/3* KD, *Ezl1* KD and *PGM* KD. G Transposons retained in EV, *PtGtsf1* KD, *Dcl2/3* KD and *Ezl1* KD. H Expression of transposons in Late, Late++ and Veg stages of EV and *PtGtsf1* KD. I Distribution of IES retention scores in EV, *PtGtsf1* KD, *Dcl2/3* KD and *Ezl1* KD. The break on the y axis indicates that the axis is discontinuous.

### The function of PtGTSF1 in transposon silencing is conserved

GTSF1 is an essential component of the piRNA pathway in animals and plays a crucial role in the repression of TEs (Arif *et al*, 2022; Chen *et al*, 2020; Dönertas *et al*, 2013; Ohtani *et al*, 2013; Yoshimura *et al*, 2018). In ciliates, the repression of TEs is achieved through DNA elimination during sexual development, and we therefore investigated whether defects in DNA elimination are the cause of the lethality observed in the absence of PtGTSF1. To do this, we isolated new MACs and extracted genomic DNA from post-autogamous cells in EV control and *PtGtsf1* KD cultures, as previously described (Arnaiz *et al*, 2012), followed by next-generation sequencing. We found the elimination of TEs belonging to LINE, SINE, SOLO-ORF and TIR families to be impaired by *PtGtsf1* KD, albeit to a weaker extent than *Dcl2/3* and *Ezl1* KD (Fig 2G). This was also accompanied with an increase in RNA expression of the corresponding TE families (Fig 2H).

Silencing of *PtGtsf1* also affected the elimination of ancient transposon remnants known as IESs. Overall, 2,783 IESs (IES retention score, IRS >= 0.1) were retained in the new MAC genome in the absence of PtGTSF1, the majority of them shorter than 200 bp (Fig 2I and EV2A, B). Nearly all the retained IESs (2,740, 98.5%) in *PtGtsf1* KD are a subset of the IESs affected by *Ezl1* KD, and two thirds (1,727) are shared with *Dcl2/3* KD (Fig 2F). Similar to the effect on TEs, the influence on IESs appears subtle, only occupying 6.2% of the total IES pool (44,817). However, as previously mentioned, *PtGtsf1* KD cells had significantly higher numbers of old MAC fragments than the control, suggesting that the degradation of the old MAC is affected (Fig 2A and B). Since this increases the contamination of old MAC DNA in *PtGtsf1* KD but not the control, the impact on TEs and IESs is likely underestimated.

To get a comprehensive view of the relationship between *PtGtsf1* and other known genes involved in IES elimination, we generated a correlation plot using IES retention scores (Fig EV2D). Although moderate, the strongest correlation is with *Rnf1* (Fig EV2D). Rnf1, also called RF4, is an accessory protein of the PRC2 complex that has been reported to act as a link between PRC2 and Ptiwi09 (Miró-Pina *et al*, 2022; Wang *et al*, 2022). Rnf1 was also among the top co-immunoprecipitated proteins in the PtGTSF1-immunoprecipitate (Fig EV1F). However, *Rnf1* KD not only impacts a larger subset of IESs than *PtGtsf1* KD (11,588 vs. 2,783), but their phenotypes differ as well (Fig EV2C). While the knockdown of *Rnf1* led to the disappearance of H3K27me3 in old MAC, this was not the case for *PtGtsf1* KD (discussed below). Therefore, the significance of their interaction warrants further investigation.

### PtGTSF1 is involved in the degradation of PIWI-bound small RNAs

Our results thus far are consistent with a conserved role of PtGTSF1 in transposon repression, as has been demonstrated for GTSF1 in metazoans. However, the piRNA-like pathway found in *P. tetraurelia* differs from the metazoan piRNA pathway in many ways, and the role of PtGTSF1 is still unclear. Additionally, the precise role of GTSF1 complexed with nuclear PIWI proteins has not yet been demonstrated in any organism. We therefore sought to determine which step of the process is affected by the absence of PtGTSF1 by interrogating the small RNA (sRNA) population. We extracted and deep sequenced sRNAs from control and *PtGtsf1* KD cells. In the control, the percentage of 25 nt scnRNAs decreased from Early to Late stages, corresponding to the degradation of self-matching MDS-scnRNAs in the old MAC (Fig 3A and B); however, this decrease did not take place in *PtGtsf1* KD (Fig 3E and F).

**Figure 3.**
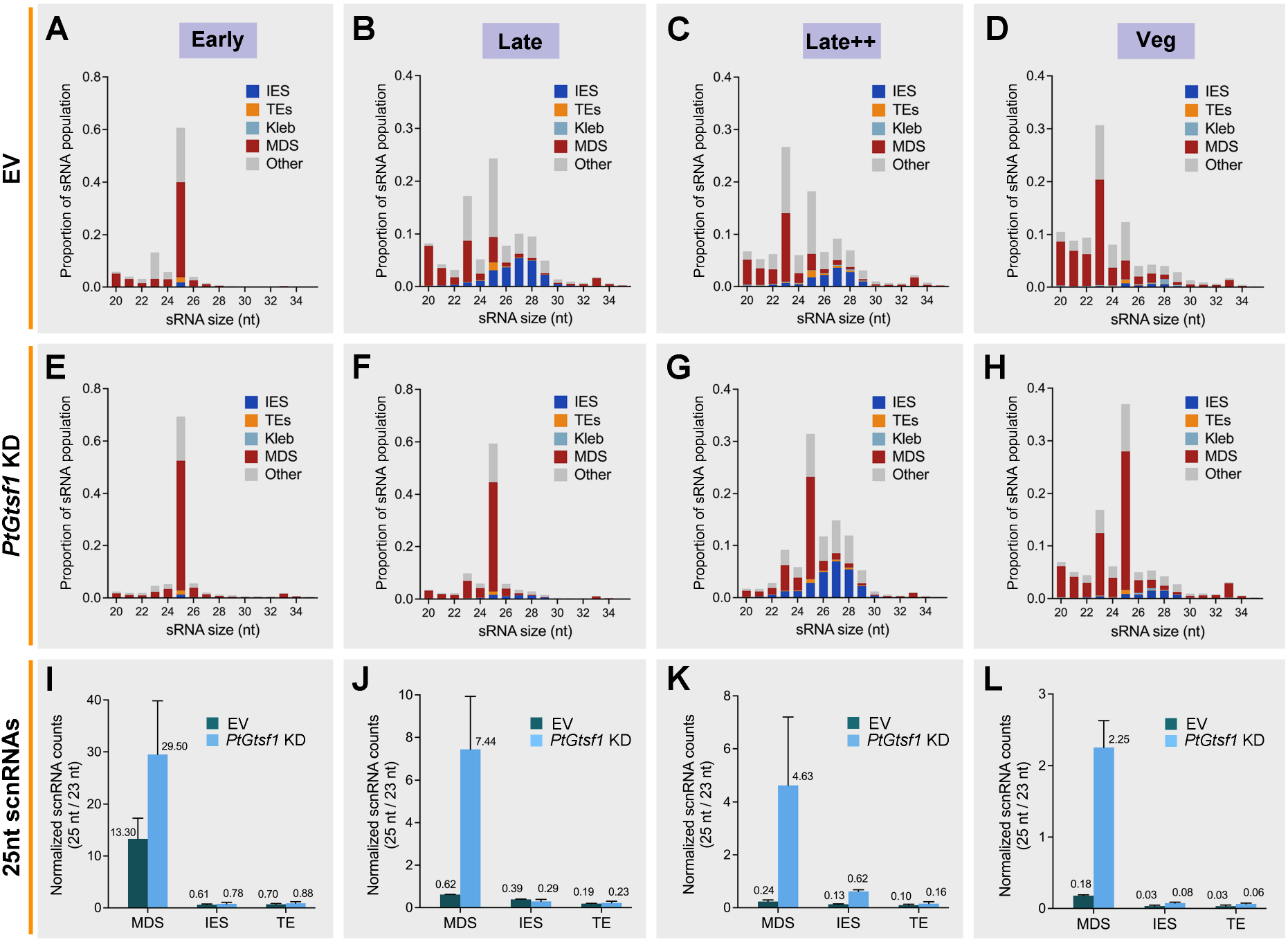
PtGTSF1 is involved in the degradation of PIWI-bound small RNAs. A, D, G, J The proportion of sRNAs mapping to various features in EV from Early to Veg stages. B, E, H, K The proportion of sRNAs mapping to various features in *PtGtsf1* KD from Early to Veg stages. C, F, I, L Normalized counts of scnRNAs mapped to MDSs, IESs and TEs in EV and *PtGtsf1* KD.

To exclude the possibility that the lower percentage of MDS-scnRNAs in the Late timepoint of the control is due to the massive production of iesRNAs, a secondary class of sRNAs ranging from 26 to 31 nt, we normalized the counts to 23 nt siRNAs. After normalization, we found the amount of MDS-scnRNAs in the Late timepoint of *PtGtsf1* KD to be much higher than that in the control, and this was also the case in Late++ and Veg stages (Fig 3J-L). Already in the Early stage, the amount of scnRNAs in *PtGtsf1* KD was higher than that in the control (Fig 3I). This may be due to an increased production of scnRNAs or because scanning already started earlier than this timepoint and MDS-scnRNAs cannot be degraded. In *P. tetraurelia*, scnRNAs are produced from the entire MIC genome, which include MDS, IES and TE sequences; however, only the amount of MDS-scnRNAs increased in *PtGtsf1* KD, while scnRNAs containing IES and TE sequences were comparable to the control (Fig 3I). Considering that PtGTSF1 does not enter MICs (Fig 1E), this is likely a problem of scnRNA degradation rather than production.

Loss of PtGTSF1 also affected another class of sRNAs involved in DNA elimination, the iesRNAs (Sandoval *et al*, 2014). In the control, iesRNAs are massively generated in the Late stage of development and gradually degrade after the elimination of IESs and TEs (Fig 3B-D). In the absence of PtGTSF1, this process is significantly delayed, and iesRNAs cannot be produced until the Late++ stage, 36 hours later than the Late stage (Fig 3F and G). While the production of iesRNAs is delayed in *PtGtsf1* KD, their degradation does not appear to be affected as they were almost entirely degraded by the Veg stage (Fig 3H). Taken together, PtGTSF1 is involved in the genome scanning process in *P. tetraurelia* by regulating PIWI-bound sRNAs, but its effects on scnRNAs and iesRNAs differ. PtGTSF1 is required for the degradation but not the production of scnRNAs and in contrast, it is needed for the duly production rather than the degradation of iesRNAs.

### Self-matching scnRNAs can enter new MACs and guide PRC2 in the absence of PtGTSF1

Since PtGTSF1 is required for scnRNA degradation and interacts with the scnRNA-binding PIWI protein Ptiwi09, we next sought to assess the location of these undegraded MDS-scnRNAs. In *P. tetraurelia*, scnRNAs bound to Ptiwi09 shuttle between different nuclei, and scnRNAs not complexed with PIWI were found to be unstable (Bouhouche *et al*, 2011; Furrer *et al*, 2017). Thus, we can visualize the location of scnRNAs using a GFP-fused Ptiwi09. Upon *PtGtsf1* KD, the entire pool of Ptiwi09-GFP appears to have entered the new MACs by the Late stage, suggesting that undegraded MDS-scnRNAs can be transferred into the new MACs (Fig 4A). Moreover, the signal of Ptiwi09-GFP in new MACs of *PtGtsf1* KD was stronger than in the control, and remained detectable even in the Veg stage (Fig 4A and B). These findings align well with our sRNA-seq results, supporting the conclusion that MDS-scnRNAs cannot be degraded without PtGTSF1 (Fig 3). Surprisingly, however, these undegraded MDS-scnRNAs appears to have entered the new MACs.

**Figure 4.**
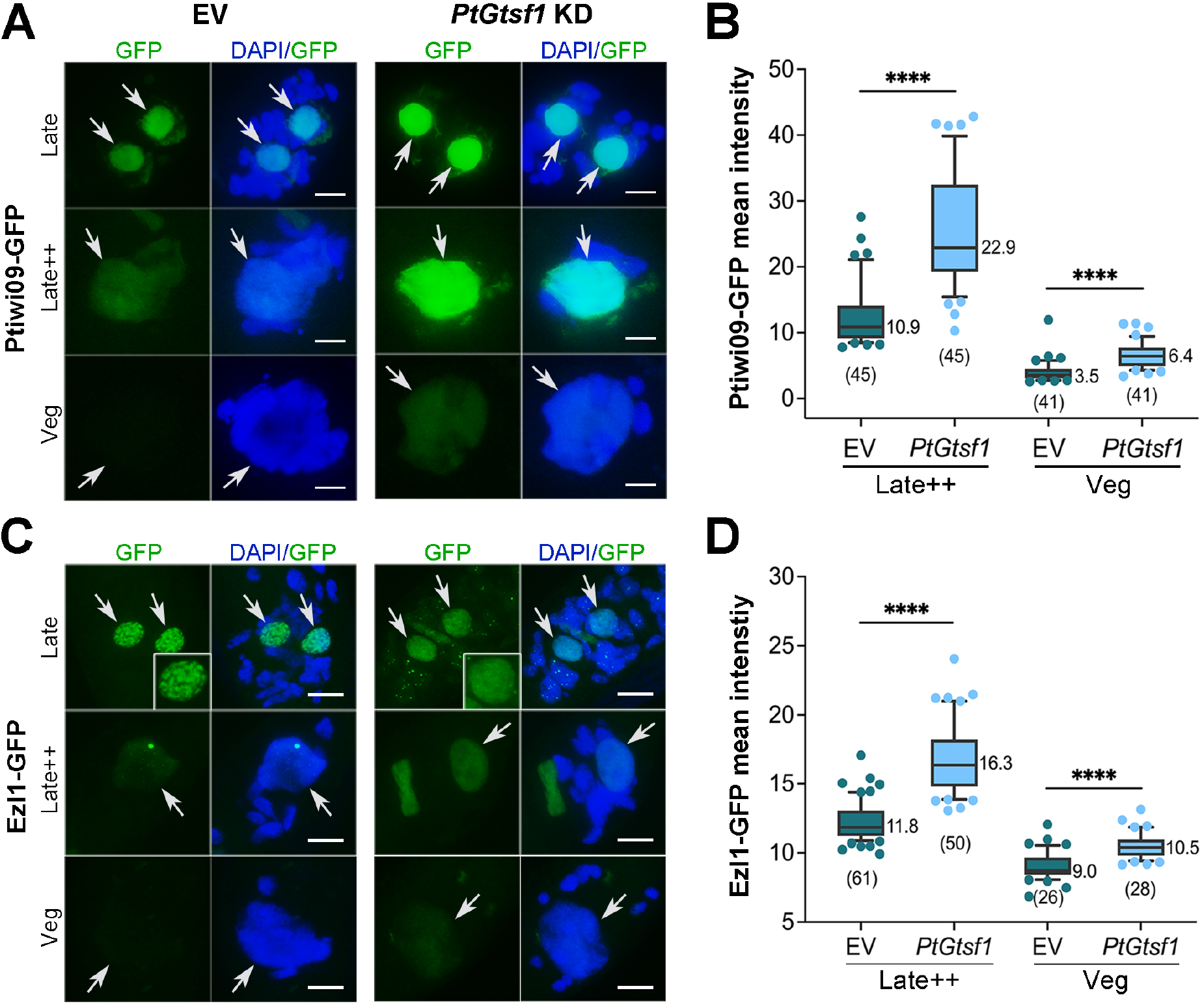
Self-matching scnRNAs can enter new MACs and guide PRC2 in the absence of PtGTSF1. A Localization of Ptiwi09-GFP in EV and *PtGtsf1* KD. Arrows indicate new MACs. Scale bar: 10μm. B Mean intensity of Ptiwi09-GFP in new MACs measured by ImageJ. Numbers in brackets under the boxes are the numbers of individuals measured. The median is listed near its line. **** *p* < 0.0001 (unpaired t-test). C Localization of Ezl1-GFP in EV and *PtGtsf1* KD. D Mean intensity of Ezl1-GFP in new MACs.

In the new MACs, scnRNAs normally guide the PRC2 complex to catalyze H3K9me3 and H3K27me3 on sequences destined for elimination (Lhuillier-Akakpo *et al*, 2014). Since the loss of PtGTSF1 affected the scnRNA population but Ptiwi09 was still able to enter the new MACs, we wondered whether these undegraded MDS-scnRNAs were “functional” (*i.e.* able to guide PRC2). We therefore investigated the localization of PRC2 in the absence of PtGTSF1 using a GFP-tagged Ezl1. In the control, the entire pool of Ezl1-GFP entered the new MACs by the Late stage of development; however, when PtGTSF1 was absent, we observed many small PRC2 foci remaining in the fragmented old MAC in the late stage (Fig 4C). Moreover, while the complex was able to enter the new MACs, it did not form foci as the control.

Since the PRC2 complex can enter new MACs but not form foci, we next assessed whether it was catalytically active by immunofluorescence of H3K9me3 and H3K27me3. In *P. tetraurelia*, both H3K9me3 and H3K27me3 are found in the new MACs in the late stage of development, and H3K27me3 is found in the old MAC in early stages as well (Lhuillier-Akakpo *et al*, 2014). However, there have been conflicting reports regarding the presence of H3K9me3 in the old MAC. One study reported that it does not exist in the old MAC at all (Lhuillier-Akakpo *et al*, 2014), whereas a second study reported detecting a faint signal in the old MAC in the early stages of autogamy (Ignarski *et al*, 2014). In our hands, we also observed it in the fragmented old MAC of wild type and two separate replicates of EV silenced cells, although the signal was weak (Figs 5A and EV3A). Of note, the localization of PRC2 complex subunits to the old MAC in the early stages of autogamy are also in agreement with these findings (Lhuillier-Akakpo *et al*, 2014). Hence, we conclude that H3K9me3 also exists in the fragmented old MAC of *P. tetraurelia*, albeit in small amounts.

**Figure 5.**
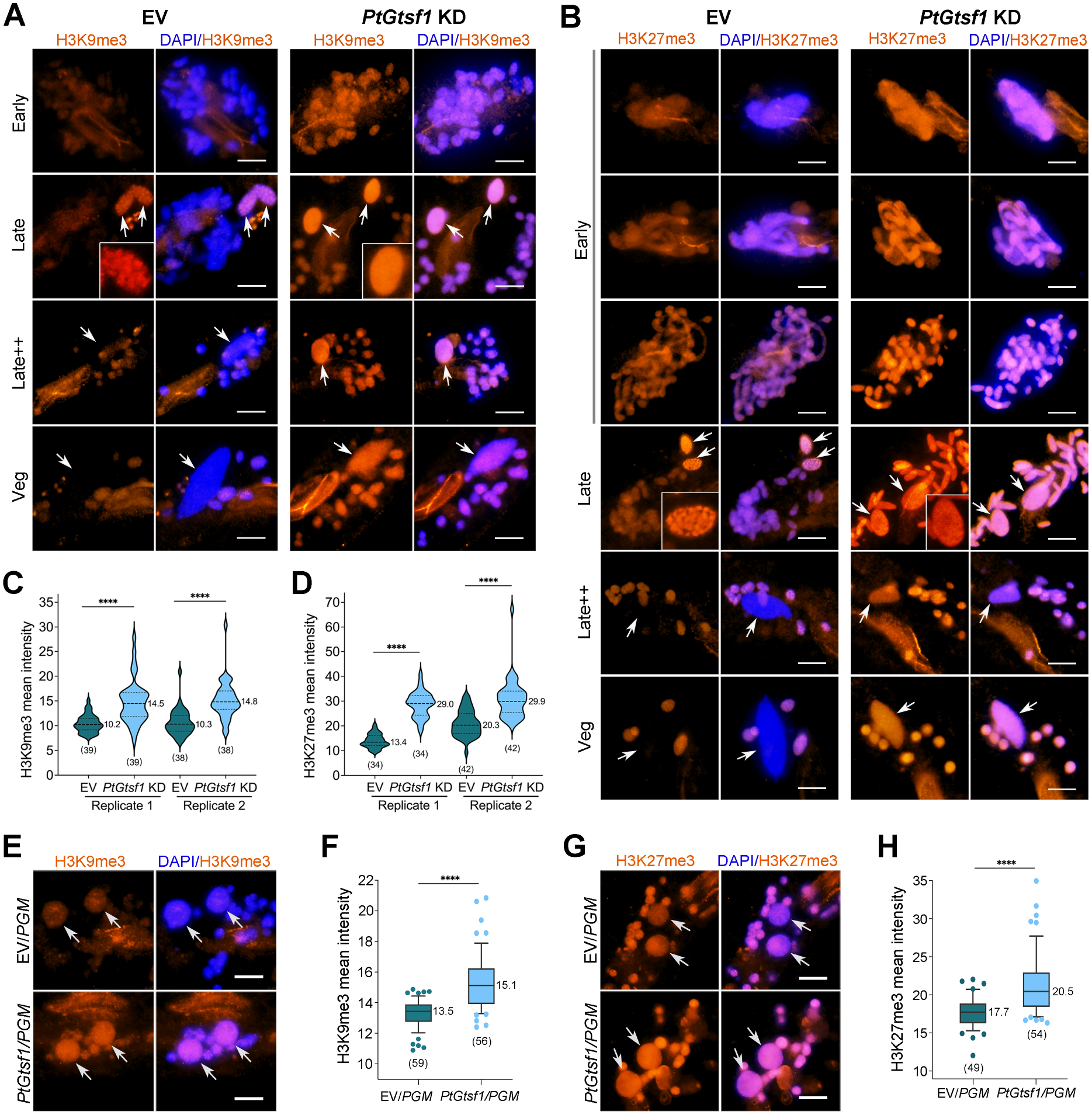
Loss of PtGTSF1 affects the intensity and distribution of H3K9me3 and H3K27me3. A Immunofluorescence of H3K9me3 during autogamy. Arrows indicate new MACs. The inset shows one of the new MACs. Scale bar: 10μm. B Immunofluorescence of H3K27me3 during autogamy. C Mean intensity of H3K9me3 in the fragmented old MAC in the Early stage. Numbers in brackets under the boxes are the numbers of individuals measured. Broken lines within the boxes are the 25% percentile, median and 75% percentile, respectively. The median is listed near its line. **** *p* < 0.0001 (unpaired t-test). D Mean intensity of H3K27me3 in the fragmented old MAC in the Early stage. E Immunofluorescence of H3K9me3 in *EV*/*PGM* KD and *PtGtsf1*/*PGM* KD. F Mean intensity of H3K9me3 in new MACs of *EV*/*PGM* and *PtGtsf1*/*PGM* KD cells. G Immunofluorescence of H3K27me3 in *EV*/*PGM* KD and *PtGtsf1*/*PGM* KD. H Mean intensity of H3K27me3 in new MACs of *EV*/*PGM* and *PtGtsf1*/*PGM* KD cells.

In early stages of both EV and *PtGtsf1* KD, H3K9me3 and H3K27me3 became detectable when the old MAC fragmented, and surprisingly, the signal in *PtGtsf1* KD was stronger than in the control (Fig 5A and B). To verify that this phenomenon was not a coincidence, we repeated the experiment and observed similar results. We also measured the mean signal intensities of both modifications in the fragmented old MAC by ImageJ, revealing that the signal intensities in *PtGtsf1* KD are significantly higher than those in the EV control (Fig 5C and D). As previously described, silencing of *PtGtsf1* did not affect scnRNA biogenesis and the MDS-scnRNAs in the control and *PtGtsf1* KD should be similar, suggesting that the increase of H3K9me3 and H3K27me3 in the old MAC is due to undegraded MDS-scnRNAs. Taken together, MDS-scnRNAs can enter new MACs and guide the PRC2 complex to catalyze H3K9me3 and H3K27me3 in the absence of PtGTSF1.

### Loss of PtGTSF1 affects the intensity and distribution of H3K9me3 and H3K27me3

The distribution of H3K9me3 and H3K27me3 in new MACs was also affected by *PtGtsf1* KD (Fig 5A and B). Normally, H3K9me3 and H3K27me3 form foci in the new MACs, which become fewer and larger as development progresses, and finally disappear from the new MACs altogether (Lhuillier-Akakpo *et al*, 2014). The formation of these foci is due to an enrichment of H3K9me3 and H3K27me3 on specific regions of the genome, such as TEs. In contrast, *PtGtsf1* KD cells had a uniform distribution of H3K9me3 and H3K27me3 in the developing new MACs, showing a consistent distribution pattern with the localization of the PRC2 complex (Fig 4C). This suggests that they are no longer confined to specific regions and are instead present throughout the entire genome. Since PRC2 can catalyze the formation of H3K9me3 and H3K27me3, this dispersed distribution is not due to an inability to bind to chromatin. We hypothesize that in the wild type situation, only scnRNAs corresponding to TE and IES sequences enter the new MACs to guide PRC2. Hence, PRC2 locates on specific regions and form polycomb foci. In the absence of PtGTSF1, scnRNAs corresponding to MDS sequences also enter new MACs and guide PRC2, causing PRC2 to locate on all kind of sequences and display a diffused distribution pattern.

Moreover, these histone modifications persisted in the new MACs of *PtGtsf1* KD cells much beyond that of the control (Fig 5A and B). By the Late++ stage, both modifications have disappeared from the new MACs in the control, whereas in *PtGtsf1* KD, they are still detectable even at the Veg stage (Fig 5A and B). This phenotype could be the result of increased methylations or alternatively, by an inability to remove these marks. If PtGTSF1 is involved in demethylation, its overexpression should result in opposite effects. Contrary to this hypothesis, the overexpression of PtGTSF1 did not affect the intensity of H3K9me3 or H3K27me3 (Fig EV3A-F). Their distribution was also not affected, and post-autogamous cells were able to return to vegetative growth, indicating that all processes of autogamy were successfully completed (Fig EV3G). Hence, the loss of PtGTSF1, but not its overexpression, affects the level and distribution of H3K9me3 and H3K27me3 in *P. tetraurelia.* We conclude that PtGTSF1 is unlikely to be involved in demethylation.

### Increased methylation levels are a result of undegraded scnRNAs

Our results suggest that PtGTSF1 is involved in the deposition rather than the removal of H3K9me3 and H3K27me3, by regulating the scnRNA-population that can guide the PRC2 complex. Moreover, the duration of both scnRNAs/Ptiwi09 and PRC2 in the new MACs were extended by *PtGtsf1* KD (Fig 4A-D), supporting the hypothesis that increased amounts of H3K9me3 and H3K27me3 are deposited in the new MACs. We further tested this hypothesis by comparing H3K9me3 and H3K27me3 levels in new MACs of control and *PtGtsf1* KD cells. However, the amount of H3K9me3 and H3K27me3 gradually decreases throughout new MAC development, making a direct comparison between control and *PtGtsf1* KD difficult. To tackle this problem, we simultaneously silenced the domesticated PiggyBac transposase PiggyMac (PGM) to retain all DNA containing H3K9me3 and H3K27me3 in both conditions. PGM is responsible for the excision of TEs and IESs in *P. tetraurelia* (Baudry *et al*, 2009; Bischerour *et al*, 2018). Even when DNA elimination is blocked, the intensity of H3K9me3 and H3K27me3 in *PtGtsf1/PGM* KD is higher than in EV/*PGM* KD (Fig 5E-H), suggesting that MDS-scnRNAs can guide the PRC2 complex to set more methylations in *PtGtsf1* KD. Hence, the increase in H3K9me3 and H3K27me3 levels are likely a result of undegraded scnRNAs.

### The localization of PtGTSF1 is independent of Ptiwi01/09 and PRC2

Although GTSF1 depletion did not impact the nuclear import of Ptiwi09, it led in an accumulation of Ptiwi09 in the new MACs by impairing the degradation of scnRNAs (Fig 4A). In other organisms, it has been reported that the localization of GTSF1 depends on PIWI (Dönertas *et al*, 2013; Yoshimura *et al*, 2018). To determine if this is also the case for PtGTSF1 in *P. tetraurelia*, we assessed the localization of PtGTSF1 when the scnRNA-binding PIWI proteins Ptiwi01 and Ptiwi09 were mislocated by silencing the Dicer-like proteins Dcl2/3. In the absence of Dcl2/3, scnRNAs cannot be produced and Ptiwi09 cannot enter the old MAC; however, this did not affect the localization of PtGTSF1 (Fig EV4A and B). Since PtGTSF1 also interacts with PRC2 and affects its localization, we assessed whether loss of PRC2, achieved by *Ezl1* KD, affected the localization of PtGTSF1. Still, the localization of PtGTSF1 remained unchanged (Fig EV4A and C). We therefore conclude that the localization of PtGTSF1 does not depend on Ptiwi01/09 nor PRC2.

### Loss of PtGTSF1 increases DNA damage in the new MACs

Depletion of PtGTSF1 increased the levels of H3K9me3 and H3K27me3, both of which are associated with gene expression regulation. We therefore assessed whether this increase also altered gene expression by mRNA-seq. In the Early and Late timepoints, only 10 and 3 genes, respectively, were differentially expressed (DEGs) (Fig 6A and B). The Late++ timepoint had 984 DEGs with 837 (85.06%) of them upregulated, while 708 DEGs (415, 58.62% upregulated) were found in the Veg timepoint (Figs 6C, D and EV5A, B). DEGs in *PtGtsf1* KD include both genes that are differentially expressed during sexual development and genes that are not (Fig EV5C-F). Notably, all genes that are known to play a role in the genome rearrangement process and are differentially expressed upon *PtGtsf1* KD are upregulated (Fig EV5G). Furthermore, neither *Ptiwi09* nor *Ezl1* are upregulated in the *PtGtsf1* KD; hence, the stronger signals of Ptiwi09 and Ezl1 in Late++ and Veg stages of *PtGtsf1* KD cannot be attributed to increased mRNA levels (Fig EV5G).

**Figure 6.**
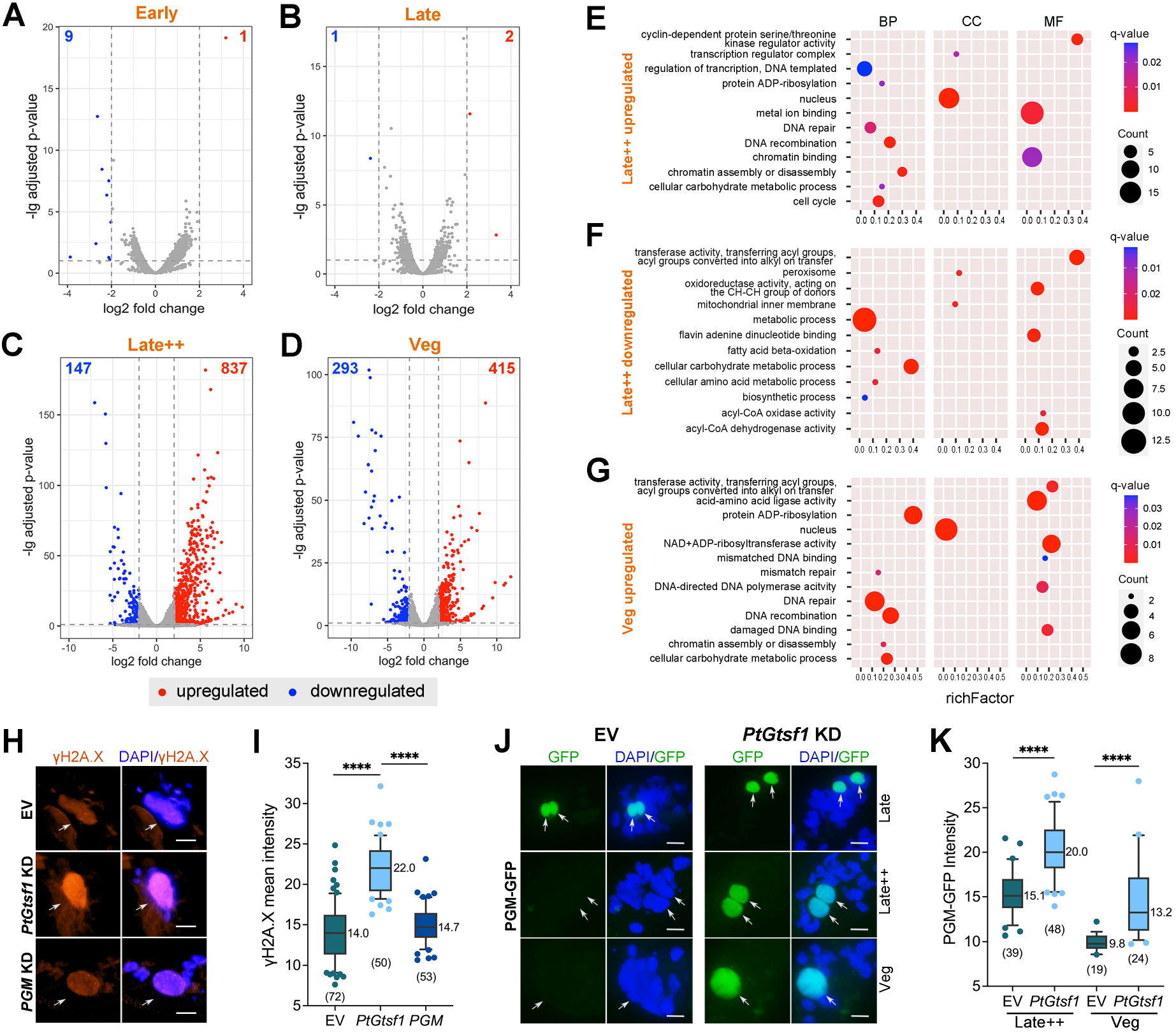
Loss of PtGTSF1 increases DNA damage in the new MACs. A-D Differentially expressed genes (DEGs) between *PtGtsf1* KD and EV in Early, Late, Late++ and Veg stages. E-G GO annotations of DEGs. Rich factor is the ratio of DEG gene numbers to the total number of genes in a certain GO term. H Immunofluorescence of γH2A.X in EV, *PtGtsf1* and *PGM* KD. Arrows indicate new MACs. Scale bar: 10μm. I Mean intensity of γH2A.X in new MACs of EV, *PtGtsf1* and *PGM* KD. Numbers in brackets under the boxes are the numbers of individuals measured. The median is listed near its line. **** *p* < 0.0001 (unpaired t-test). J Localization of PGM-GFP in EV and *PtGtsf1* KD. K Mean intensity of PGM-GFP in new MACs of EV and *PtGtsf1* KD.

We further predicted the functions of the DEGs in Late++ and Veg timepoints by GO analysis (Fig 6E-G). Many upregulated genes are related to DNA damage and repair, such as DNA repair, DNA recombination, mismatch repair, mismatched DNA binding, and damaged DNA binding, suggesting that there may be more DNA damage occurring when *PtGtsf1* is silenced. To investigate this, we assessed the levels of DNA damage by immunofluorescence of γ-H2A.X (Fig 6H). Since DNA elimination by default massively induces DNA double-strand breaks (DSBs), we first investigated the localization of the excision complex using a GFP-fused PGM. In the wild type situation, PGM disappears after the Late++ stage, when DNA elimination is completed (Fig 6J and K) (Baudry *et al*, 2009). Interestingly, PGM was still present even in Veg stages in the *PtGtsf1* KD, mirroring our findings for Ptiwi09 and the PRC2 complex (Figs 4 and 6J, K). We next investigated whether blocking DNA elimination by *PGM* KD affects DNA damage levels in the new MACs. Silencing *PGM* effectively blocks DNA elimination but did not lead to lower DNA damage levels compared to the EV control, suggesting that DSBs caused by DNA elimination are quickly repaired (Fig 6H and I). This is likely due to efficient coupling of DNA elimination and repair, as previously reported (Marmignon *et al*, 2014). In contrast, *PtGtsf1* KD cells had significantly higher levels of DNA damage than the EV and *PGM* KD (Fig 6H and I). This suggest that more DSBs are occurring in the absence of PtGTSF1, which may reflect additional cleavage events caused by MDS-scnRNAs and increased methylation marks in the new MACs. Alternatively, the absence of PtGTSF1 could affect DNA repair after cleavage. Taken together, our findings suggests that the increase in DNA damage in *PtGsf1* KD may be the result of increased activity of the excision complex.

## Discussion

Findings in several organisms have established GTSF1 as an essential component of the piRNA pathway in animals. Here, we expanded on these findings by revealing that the PIWI-GTSF1 interaction also exists in the unicellular eukaryote *P. tetraurelia*, with the conserved function of TE silencing. In *P. tetraurelia*, PtGTSF1 is essential for sexual development by facilitating the degradation of small RNAs recognizing the organism’s own genomic sequences, as a way to distinguish self-from non-self-DNA.

### PtGTSF1 is required for the elimination of Ptiwi01/09 after scnRNA-target RNA pairing

Previous studies have demonstrated that the PRC2 complex requires scnRNAs to set H3K9me3 and H3K27me3 on TEs, which aids in their recognition (Lhuillier-Akakpo *et al*, 2014). In the absence of PtGTSF1, scnRNAs containing self-matching sequences cannot be removed, but we found that they are still able to guide the PRC2 complex to form H3K9me3 and H3K27me3 in both the old and new MACs (Figs 3 and 5). Since the PRC2 complex requires scnRNAs to deposit these marks, pairing between scnRNAs and the nascent transcripts of the old MAC still takes place in the absence of PtGTSF1, which places PtGTSF1 downstream of scnRNA-target RNA pairing. Interestingly, this appears to be a PtGTSF1-specific phenotype, as this does not occur in known cases of scnRNA-degradation defects. In our previous work, we constructed a “Late-PRC2” strain of *P. tetraurelia* whose PRC2 complex only locates and acts in the new MACs (Wang *et al*, 2022). Loss of PRC2 in the old MAC impaired the degradation of MDS-scnRNAs similar to what we found for *PtGtsf1* KD in this study. These undegraded MDS-scnRNAs led to a diffused distribution of PRC2 in the new MACs; however, and in contrast to the case in *PtGtsf1* KD, H3K9me3 and H3K27me3 still formed foci as the control. These results suggest that the characteristics of scnRNA-Ptiwi09 complexes before and after pairing differ. Thus, we speculate that scnRNA-Ptiwi09 complexes are marked after pairing with its targets, and that this process depends on PtGTSF1. This mark, on scnRNAs or Ptiwi09, would then result in the degradation of the scnRNA-Ptiwi09 complexes.

The elimination of successfully paired scnRNAs is of outmost importance as this allows the organism to distinguish self from non-self-DNA. In the absence of such selection, essential DNA sequences might be deleted; alternatively, the retention of IESs would disrupt gene expression in the new MACs. Hence, the organism requires an efficient degradation system that can distinguish paired and unpaired scnRNAs. While the details of this process are still unknown, one can imagine that this may occur either at the sRNA level or at the protein level. Interestingly, previous studies found that Argonaute carrying an extensively paired miRNA triggers its degradation through ubiquitination in a process known as target-directed miRNA degradation (TDMD) (Han *et al*, 2020; Shi *et al*, 2020). Considering that Ptiwi09-bound scnRNAs have a perfect match with its targets, we envision that a similar mechanism may be at play in *P. tetraurelia*, in which Ptiwi09 proteins carrying perfectly matched scnRNAs undergo a conformational change that results in their degradation, in a process dependent on PtGTSF1.

Our study provided insight into the mechanisms governing the genome scanning process, summarized in the following model (Fig. 7): At the beginning of autogamy, MICs are transcribed to generate scnRNAs corresponding to MDSs, IESs and TEs, all of which bind with Ptiwi01/09 and enter the old MAC to identify self-matching MDS-scnRNAs by pairing with transcripts originating from the old MAC genome. After pairing, MDS-scnRNAs guide PRC2 to form H3K9me3 and H3K27me3 in the old MAC, after which MDS-scnRNAs are degraded. The unpaired IES- and TE-scnRNAs are transferred into the new MACs to guide the elimination of IESs and TEs.

**Figure 7.**
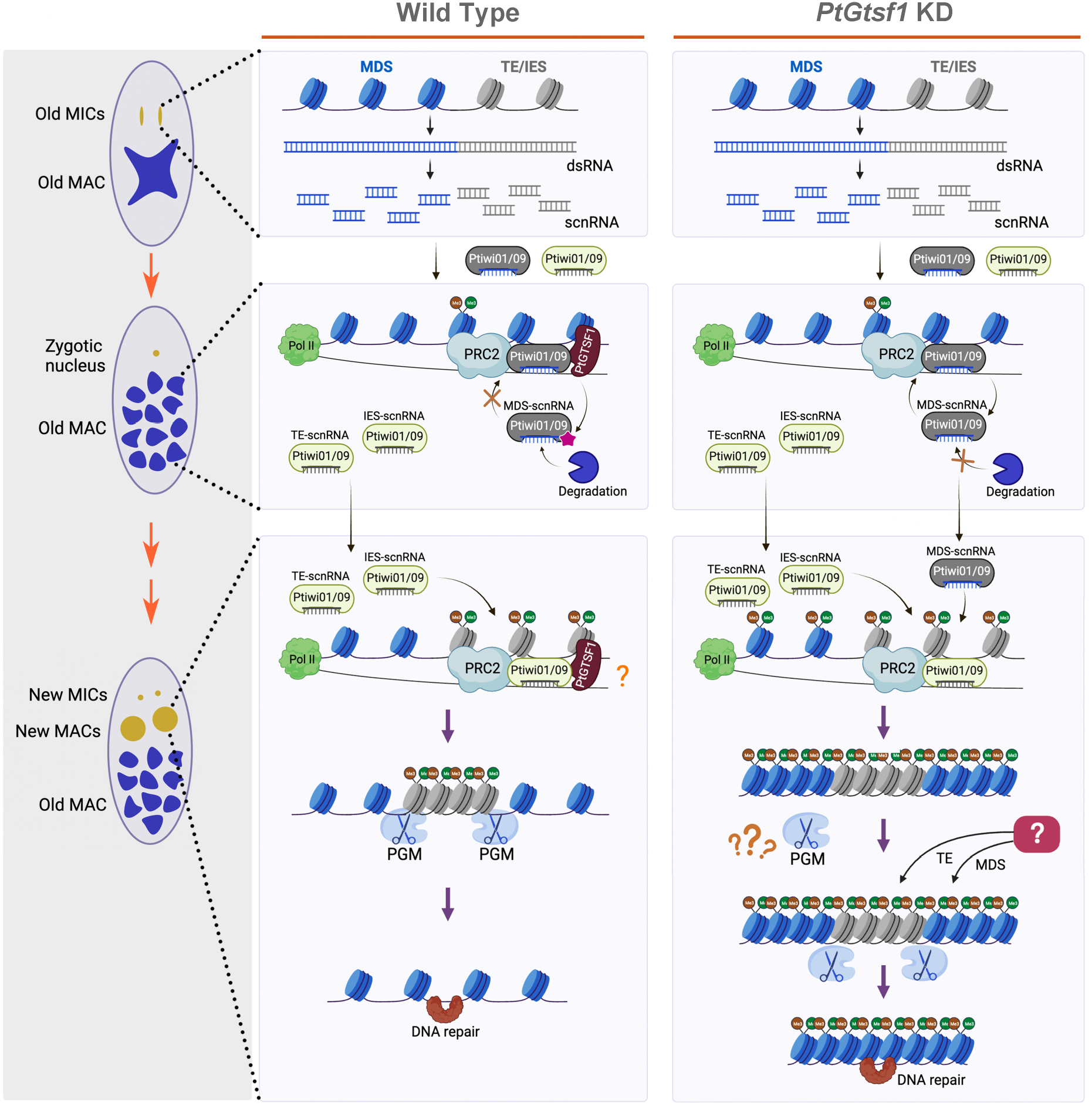
Model depicting the role of PtGTSF1 in the genome rearrangement process of *Paramecium tetraurelia.* During autogamy, micronuclei (MICs) are bidirectionally transcribed to produce long ncRNAs that are processed into scnRNAs, which then bind with Ptiwi01/09 and enter the old macronucleus (MAC). In the old MAC, scnRNAs pair with nascent transcripts generated from the old MAC genome. Paired scnRNAs (MDS-scnRNAs) guide the PRC2 complex to set H3K9me3 and H3K27me3, and are then marked in a PtGTSF1-dependent manner, resulting in their degradation. Unpaired scnRNAs enter the new MACs to identify transposons (TEs) and internally eliminated sequences (IESs). Targeting by scnRNAs guide the PRC2 complex to set H3K9me3 and H3K27me3 on TEs, resulting in heterochromatin formation and recruitment of PiggyMAC (PGM) and related proteins. PGM then cleaves and eliminates the TEs. When PtGTSF1 is absent, MDS-scnRNAs cannot be degraded and aberrantly enter the new MACs, resulting in an abnormal distribution of H3K9me3 and H3K27me3, thereby affecting the recognition of TEs. Additional mechanisms yet to be discovered likely help recognize TEs. Figure created with BioRender.com.

### The possibility of protective marks in *P. tetraurelia*

The enrichment of H3K9me3 and H3K27me3 on TE sequences is essential for their elimination (Frapporti *et al*, 2019). These modifications facilitate the heterochromatinization of TEs regions, which are subsequently recognized by PGM and related proteins (Frapporti *et al*, 2019; Wang *et al*, 2022). PGM is then believed to cleave the boundaries of heterochromatin and euchromatin to eliminate the TEs. The loss of PtGTSF1 leads to the aberrant entry of MDS-scnRNAs into the new MACs, which then guides PRC2 to form H3K9me3 and H3K27me3. Despite the presence of these methylation marks, a subset of TEs are still retained in *PtGtsf1* KD. A plausible explanation for the DNA elimination defect despite the presence of H3K9me3 and H3K27me3 is that without PtGTSF1, these marks are distributed throughout the entire genome, resulting in an inability of the PGM complex to distinguish TEs by the state of the chromatin. Since most IESs and TEs are still eliminated in the new MACs, there may also be additional mechanisms to recognize scnRNAs of MDS, TE and IES origin. In the related ciliate *Tetrahymena thermophila*, the boundaries of MDSs are protected to avoid excessive elimination (Suhren *et al*, 2017). Though this may differ from the case in *P. tetraurelia*, it is possible that the MDSs might also be protected in this organism. This, in turn, may aid in the identification of TEs and IESs.

### H3K9me3 and H3K27me3 are unlikely to be repressive marks in *P. tetraurelia*

In the absence of PtGTSF1, the levels of H3K9me3 and H3K27me3 increased in the old MAC due to undegraded MDS-scnRNAs. Currently, our knowledge regarding the function of H3K9me3 and H3K27me3 in the old MAC is very limited. One function we would expect based on their roles in other organisms is gene expression regulation; however, no matter the increase or decrease of these marks in the old MAC, it appears to have a minimal impact on gene expression. Moreover, while H3K9me3 and H3K27me3 are classically considered as repressive marks, we found most genes to be upregulated when these marks increased in *P. tetraurelia.* In our previous work, we reported that the majority of DEGs in the absence of the PRC2 complex (*i.e.* loss of H3K9me3 and H3K27me3) are downregulated (Wang *et al*, 2022). Additionally, Drews *et al* (2022) reported H3K27me3 to be enriched on highly-expressed genes, all of which are consistent with our current observations. In light of multiple lines of evidence, we believe that these histone marks are not repressive in *P. tetraurelia*.

### The role of GTSF1 in regulating the activity of nuclear PIWI proteins

It is important to note that the role of GTSF1 has only been elucidated for catalytically active PIWI proteins, in which it appears to enhance the intrinsically weak RNA cleavage activities of PIWI (Arif *et al*, 2022). In mouse, silk moth and sponges, GTSF1 interacts with PIWI and recognizes the pre-catalytic piRISC state, facilitating a conformational change in PIWI and stabilizing its catalytically active conformation (Arif *et al*, 2022). Although Ptiwi09 is also a catalytically active PIWI protein, its role in the nucleus suggests that its main function is not to directly cleave its targets, but rather that it acts similarly to the catalytically inactive *Drosophila* Piwi, which guides downstream factors to deposit histone methylation marks (Brower-Toland *et al*, 2007; Le Thomas *et al*, 2013). Of note, *Drosophila* Piwi also requires GTSF1 for its activity (Dönertas *et al*, 2013; Muerdter *et al*, 2013; Ohtani *et al*, 2013). We can envision a model in which the binding of nuclear PIWI proteins to its targets induces a conformational change in PIWI, and that the interaction with GTSF1 can only happen after this change.

Our results are consistent with a conserved role of PtGTSF1 as in metazoans, in which its canonical function is transposon silencing through its interaction with, and regulation of, PIWI proteins. Protists are the ancestors of multicellular organisms; hence, the existence of GTSF1 in protists suggests that this protein appeared before multicellularity. Additionally, PtGTSF1 is also involved in transposon repression in *P. tetraurelia*, demonstrating that transposon repression is an ancient function of GTSF1.

## Materials and methods

### *Paramecium tetraurelia* cultivation and autogamy induction

*Paramecium tetraurelia* strain 51, mating type seven, was used to perform the experiments. Cells were cultured in wheat grass power (WGP) medium (Pine international, Lawrence, KS) bacterized with *Klebsiella pneumoniae* and supplied with 0.8mg/L β-sitosterol (Shanghai Yuanye Bio-technology) (Beisson *et al*, 2010b). Autogamy was induced by starvation.

### Gene silencing

Gene silencing was carried out as described in Beisson *et al* (2010c). A fragment of the coding region from 1 to 366 bp of *PtGtsf1* (Gene ID: PTET.51.1.G0490019) was cloned into the L4440 plasmid and then transformed into the HT115 (DE3) strain of *Escherichia coli*. Silencing constructs of *Ezl1*, *Dcl2*, *Dcl3* and *PGM* were made according to published papers (Baudry *et al*, 2009; Lepere *et al*, 2009; Lhuillier-Akakpo *et al*, 2014).

Briefly, silencing was performed as follows. Bacteria with silencing constructs were cultured overnight in LB medium, and then 1:100 diluted in 1x WGP medium and incubated overnight. The culture was further diluted with 1x WGP to an OD_600_ of 0.04 and allowed to grow until the OD_600_ reached a range between 0.07 and 0.1. Isopropyl-β-d-thiogalactopyranoside (IPTG) was added to a final concentration of 0.4mM and the culture was incubated for at least 4 hours to induce the production of double-stranded RNA (dsRNA). Prior to seeding, the silencing medium was supplemented with 0.8mg/L β-sitosterol and the *P. tetraurelia* cells were washed in Tris-HCl (10mM, pH 7.5) and seeded at a density of 200 cells/ml.

### DNA preparation and microinjection

The entire *PtGtsf1* gene including flanking sequences (206 bp upstream and 179 bp downstream), was cloned into the pCE2 vector using the 5min^TM^ TA/Blunt-Zero Cloning Kit (Vazyme Biotechnology). A codon-optimized GFP or FlagHA was then inserted after the start codon ATG of *PtGtsf1*. Plasmids containing GFP tagged Ptiwi09, Ezl1 and PGM were constructed as in the published papers (Baudry *et al*, 2009; Bouhouche *et al*, 2011; Lhuillier-Akakpo *et al*, 2014). After confirmation of the plasmids by sequencing, the constructs were extracted from bacteria and linearized by enzyme digestion in the backbone, followed by purification using phenol chloroform (pH 8.0) and Ultrafree-MC Centrifugal Filters (Millipore).

To perform microinjections, post-autogamous *P. tetraurelia* cells were recovered to the vegetative stage and grown for 4 to 8 divisions. DNA was injected into the macronucleus as described in (Beisson *et al*, 2010a). Successful injections were confirmed by PCR and the cells grown up for follow-up experiments.

### Immunofluorescence

Around 150,000 cells were centrifuged and washed twice with 10mM Tris-HCl (pH 7.5). The cells were permeabilized with 1% Triton X-100 in 1x PHEM buffer (10mM EGTA, 25mM HEPES, 2mM MgCl_2_, 60mM PIPES (pH 6.9)) for 10 minutes and fixed with 2% Paraformaldehyde in 1x PHEM buffer for 10 minutes. Next, the cells were blocked in 3% BSA in TBSTEM buffer (10mM EGTA, 2mM MgCl_2_, 0.15M NaCl, 10mM Tris-HCl, 1% Tween 20) for 30 minutes. After blocking, the cells were incubated with a 1:200 dilution of the following antibodies overnight at 4°C: anti-trimethyl-Histone H3 (Lys27) (07-449, Millipore), anti-trimethyl-Histone H3 (Lys9) (07-442, Millipore) or anti-gamma H2A.X (phosphor S139) antibody (ab11174, abcam). Afterwards, the cells were washed twice with 3% BSA in TBSTEM and incubated with a goat anti-rabbit Alexa Fluor 546 Secondary antibody (A-11071, Invitrogen) at a dilution of 1:4,000 for an hour. Following two washes with 1x phosphate-buffered saline (PBS), the cells were spread onto glass slides and mounted with ProLong Glass Antifade Mountant (P36980, Invitrogen), containing the blue DNA stain NucBlue. Imaging was performed with a Axio Imager D2 (Zeiss) and the Zen 2 software.

### Imaging and measurement of the mean fluorescence intensity

For cells injected with GFP-tagged constructs, approximately 150,000 cells at the desired timepoint were centrifuged and fixed in 75% ethanol. Before imaging, the cells were washed three times with 1x PBS and stained with 4,6-diamidino-2-phenylindole (DAPI). Then, the cells were spread onto glass slides for imaging. All imaging was performed with a Axio Imager D2 (Zeiss) and the Zen 2 software.

Images used to compare signal intensities were taken under the same conditions. ImageJ (Schneider *et al*, 2012) was used to measure the fluorescence intensity with background subtraction and regions of interested selected by polygon selections.

### Immunoprecipitation

Around 1.5 million cells were collected and washed twice with 10mM Tris-HCl (pH 7.5), followed by two washes with cold 1× PBS. After centrifugation and removal of the supernatant, the cell pellet was resuspended in 2 ml of lysis buffer (50mM Tris-HCl (pH 8.0), 150mM NaCl, 5mM MgCl_2_, 1mM DTT, 1× complete EDTA-free protease inhibitor cocktail tablet (Roche), 1% Triton X-100, 10% glycerol). The samples were subjected to sonication using a Branson digital sonifier SFX250, with an amplitude of 55% for 15s. Following sonication, the soluble fraction was collected after clearing by centrifugation at 15,060rpm for 30min. Next, 50μl of Anti-HA Affinity Matrix (11815016001, Roche) was washed three times with IP buffer (10mM Tris-HCl (pH 8.0), 150mM NaCl, 1mM MgCl_2_, 0.01% NP40, 5% glycerol) and the soluble fraction (1 ml) was added to the beads, followed by an overnight incubation at 4°C. After the incubation, the beads were washed six times with IP buffer, the supernatant discarded, and the beads fraction used for further experiments.

### Western blot

Immunoprecipitated proteins were separated by SDS-PAGE electrophoresis and transferred to a nitrocellulose membrane (Amersham Protran, GE Healthcare Life Sciences) using a wet transfer method. The membrane was blocked with 10% skim milk in PBST (1× PBS with 0.1% Tween-20) for 1 hour at room temperature. After blocking, the membrane was incubated overnight at 4°C with a Rabbit anti-HA primary antibody (3724S, Cell Signaling Technology) at a dilution of 1:1000. Next, the membrane was washed three times with PBST and incubated with a 1:8000 diluted secondary antibody (ProteinFind^®^ Goat Anti-Rabbit IgG (H+L), HRP Conjugate, HS101-01, Transgen Biotechnology) at room temperature for 1 hour. The membrane was washed three times with PBST, followed by one wash with PBS. Pierce^TM^ ECL Western Blotting Substrate (32106, Thermo Scientific) was used to develop the signal.

### Mass spectrometry

Immunoprecipitated proteins were separated by SDS-PAGE electrophoresis, stained with Coomassie brilliant blue R-250 (Solarbio) and cut into gel slices. After destaining and dehydration, the gel bands were subjected to reduction by incubation with 10 mM DTT at 56°C for 1 hour, followed by alkylation with 55 mM IAA for 45 min. The proteins were digested with trypsin overnight at 37°C. The resulting peptides were separated by the UltiMate3000 RSLCnano ultra-high performance liquid system using a gradient of 65 min at the rate of 400 nL/min. Solvent A was water with 0.1% FA and solvent B was 98% ACN with 0.1% FA. The gradient was 5-8% B for 6min, 8-30% B for 34min, 30-60% B for 5 min, 60-80% B for 3 min, 80% B for 8min, 80-5% B for 2 min and 5% B for 7 min. The peptides were then analyzed by the Thermo Scientific^TM^ Q Exactive^TM^ mass spectrometer in Data Dependent Acquisition (DDA) mode. The ion source voltage was 1.8 kV, the full scan range of MS was 350-2000 m/z, and the scanning resolution was set to 70,000. The top 20 peptides were selected for MS2 and the resolution was 17,500. The proteomic data was queried using Mascot (2.3.0) against the protein sequences of *Paramecium tetraurelia* downloaded from ParameciumDB <u>(ptetraurelia_mac_51_annotation_v2.0.protein.fa)</u>. The search parameters were set as follows: the enzyme was specified as trypsin; the missed cleavage was set to 2; the mass tolerance was 15 ppm for precursor and 20 mmu for fragment; the fixed modification was cysteine carbamidomethylation; the variable modifications included N-terminal glutamate to pyroglutamate (Gln → pyro-Glu) and methionine oxidation; the ion score was set above 23.

### Amino acid sequence alignment and Neighbor-Joining tree construction

The amino acid sequences of GTSF1 from different organisms were downloaded from the UniProt database with the accession numbers shown in Figure S1. The sequences were aligned in Geneious with MUSCLE alignment and the Neighbor-joining tree was constructed with Geneious Tree Builder using the genetic distance model of Jukes-Cantor.

### Survival test

After undergoing autogamy, the cells were transferred into individual wells containing bacterized 0.2× WGP (0.8mg/L of β-sitosterol added) to recover to a vegetative stage. The growth of the cells was monitored for three consecutive days. If the cells grew similarly as the control, they were categorized as healthy, otherwise as sick (fewer cells than the control) or dead (only one or no alive cell).

### Macronuclear isolation and DNA extraction

The macronuclei of post-autogamous cells were isolated following the protocol in Arnaiz *et al* (2012). Approximately 1.5 million cells were centrifuged and washed twice with 10mM Tris-HCl (pH 7.5). The cell pellet was resuspended in 2.5 volumes of lysis buffer 1 (0.25M sucrose, 10mM MgCl_2_, 10mM Tris-HCl (pH 6.8), 0.2% NP-40). After incubating on ice for 5 minutes, a Potter-Elvehjem homogenizer was used to disrupt the cell membranes while keeping the nuclei intact. Subsequently, the lysate was washed twice with wash buffer (0.25M sucrose, 10mM MgCl_2_, 10mM Tris-HCl (pH 7.5)). The pellet was resuspended in 3 volumes of sucrose buffer (2.1M sucrose, 10mM MgCl_2_, 10mM Tris-HCl (pH 7.5)), and carefully layered on top of 3ml of sucrose buffer in a centrifuge tube (344060, Beckman Coulter). The tube was filled with wash buffer to create a sucrose gradient. The macronuclei were isolated by ultracentrifugation at 35,000 rpm, 4°C for an hour using a Beckman Optima L-90K Ultracentrifuge. Following ultracentrifugation, macronuclear DNA was extracted by phenol chloroform. Next-generation sequencing was performed by the Novogene company (Tianjin, China).

### RNA extraction, mRNA-seq and sRNA-seq

Half a million cells at the desired timepoint were collected and washed twice with 10mM Tris-HCl (pH 7.5). After removing the supernatant, the cells were frozen in liquid nitrogen until RNA extraction. Total RNA was extracted with TRIzol (Sigma-Aldrich), following the TRIzol reagent BD protocol.

The RNA was sequenced by the Novogene company on a Novaseq 6000 to obtain paired-end 2x 150bp (mRNA sequencing) and single-end 1x 50bp (sRNA sequencing).

### Reference genomes

The following reference genomes were used in the IES and transposon analyses, and for read mapping:

MAC: http://paramecium.cgm.cnrs-gif.fr/download/fasta/ptetraurelia_mac_51.fa

MAC+IES: http://paramecium.cgm.cnrs-gif.fr/download/fasta/ptetraurelia_mac_51_with_ies.fa

TE: https://paramecium.i2bc.paris-saclay.fr/files/Paramecium/tetraurelia/51/annotations/ptetraurelia_mic2/ptetraurelia_TE_consensus_v1.0.fa

### sRNA-seq analysis

Single-end sRNA reads ranged from 20 to 45 nt were processed using fastp (-q 20 -l 20, v0.23.1,https://doi.org/10.1093/bioinformatics/bty560) before mapping using Bowtie (v1.3.1; (Langmead *et al*, 2009)) with default parameters. The size-selected reads were mapped to the following reference datasets sequentially: the genome of *Klebsiella* (CP002824.1); IESs from the MAC+IES genome (ptetraurelia_mac_51_with_ies.fa); the germline MAC genome (referred to as MDSs)(ptetraurelia_mac_51.fa). The 23 nt siRNA reads mapped to the RNAi targets were removed and the scnRNA read counts were normalized using the total number of reads and siRNAs.

### mRNA-seq analysis

Paired-end reads were filtered using fastp with -q 20, -l 75, -c, --detect_adapter_for_pe (v0.23.1; (Chen *et al*, 2018)), and then mapped against the MAC genome of *P. tetraurelia* (ptetraurelia_mac_51.fa) using hisat2 (v2.2.1; (Kim *et al*, 2019)) with default parameters. Reads were counted using featureCounts (v2.0.1, (Liao *et al*, 2014)) with the parameter of -p and the output used as the input for DESeq2 (Love *et al*, 2014). Differentially expressed genes in the *PtGtsf1* KD were identified as those with an adjusted p value less than 0.1 and with at least a four-fold change relative to the corresponding control timepoint. Genes were classified as upregulated (fold change ≥4) or downregulated (fold change ≤¼). Gene ontology functional enrichment was performed using the clusterProfiler package (Yu *et al*, 2012).

### IES retention scores and correlation plots

IES retention scores (IRSs) were calculated with the MIRET component of ParTIES (Denby Wilkes *et al*, 2016) using the whole score option. Correlation plots between different gene knockdowns were calculated based on IES retention scores (IRSs) using After_ParTIES (Swart *et al*, 2017).

## Data availability

The mRNA-seq, sRNA-seq and genomic DNA sequencing data were submitted to NCBI database under the BioProject PRJNA1020915.

## Acknowledgements

We thank Prof. Weibo Song from Ocean University of China for his suggestion and help in drafting the manuscript. We acknowledge the computing resources provided on IEMB-1, a high-performance computing cluster operated by the Institute of Evolution and Marine Biodiversity. This research was supported by the National Natural Science Foundation of China (32100382, 32270539, 31961123002), the Natural Science Foundation of Shandong Province (ZR2021QC104, ZR2020JQ13), Young Elite Scientists Sponsorship Program by CAST (2022QNRC001), and the Fundamental Research Funds for the Central Universities (202141004). This work was supported by the World Premier International Research Center Initiative (WPI), MEXT, Japan.

## Author Contribution

**Chundi Wang:** Conceptualization; data curation; formal analysis; investigation; visualization; writing-original draft; writing – review and editing; funding acquisition. **Liping Lv:** Formal analysis; visualization; writing – original draft. **Therese Solberg:** Conceptualization; writing-review and editing. **Zhiwei Wen:** Investigation; **Haoyue Zhang:** Investigation. **Feng Gao:** Supervision; writing – original draft; writing – review and editing; funding acquisition.

## Expanded View Figure Legends

**Figure EV1.**
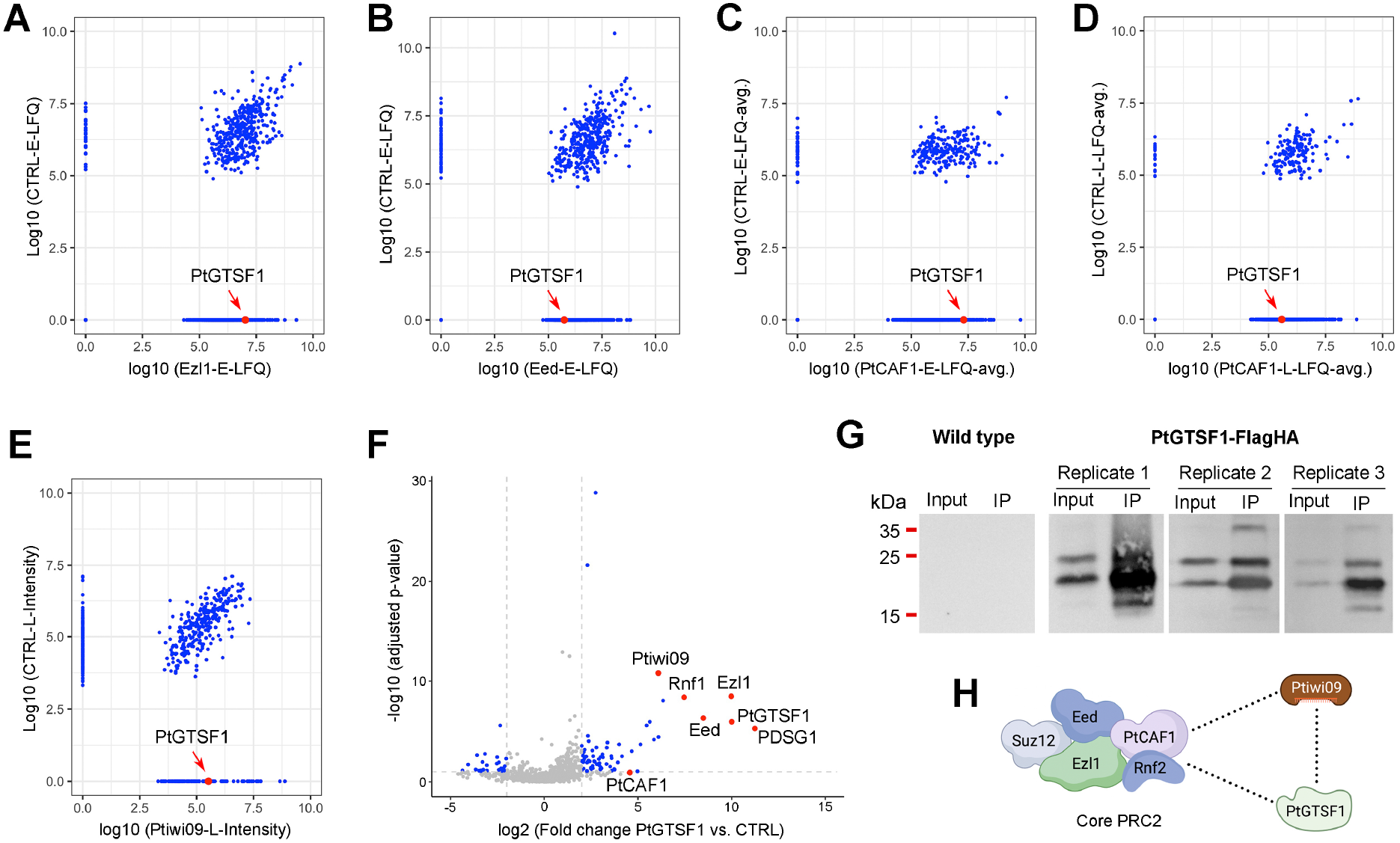
PtGTSF1 interacts with Ptiwi09 and PRC2 in *Paramecium tetraurelia*. A-E Scatter plots of detected proteins in the mass spectrometry data from Ezl1 (Early timepoint), Eed (Early timepoint), PtCAF1 (Early and Late timepoints) and Ptiwi09 (Late timepoint) immunoprecipitated proteins, with wild type of the same timepoints as controls. PtGTSF1 is indicated with a red dot and arrow. F Volcano plot of PtGTSF1 co-immunoprecipitated proteins. PRC2 complex subunits, Ptiwi09 and PDSG1 are indicated with red dots and labels. G Western blots of wild type and PtGTSF1-FlagHA immunoprecipitate using an anti-HA antibody to detect PtGTSF1. H Schematic to show the interaction between PtGTSF1, Ptiwi09 and PRC2.

**Figure EV2.**
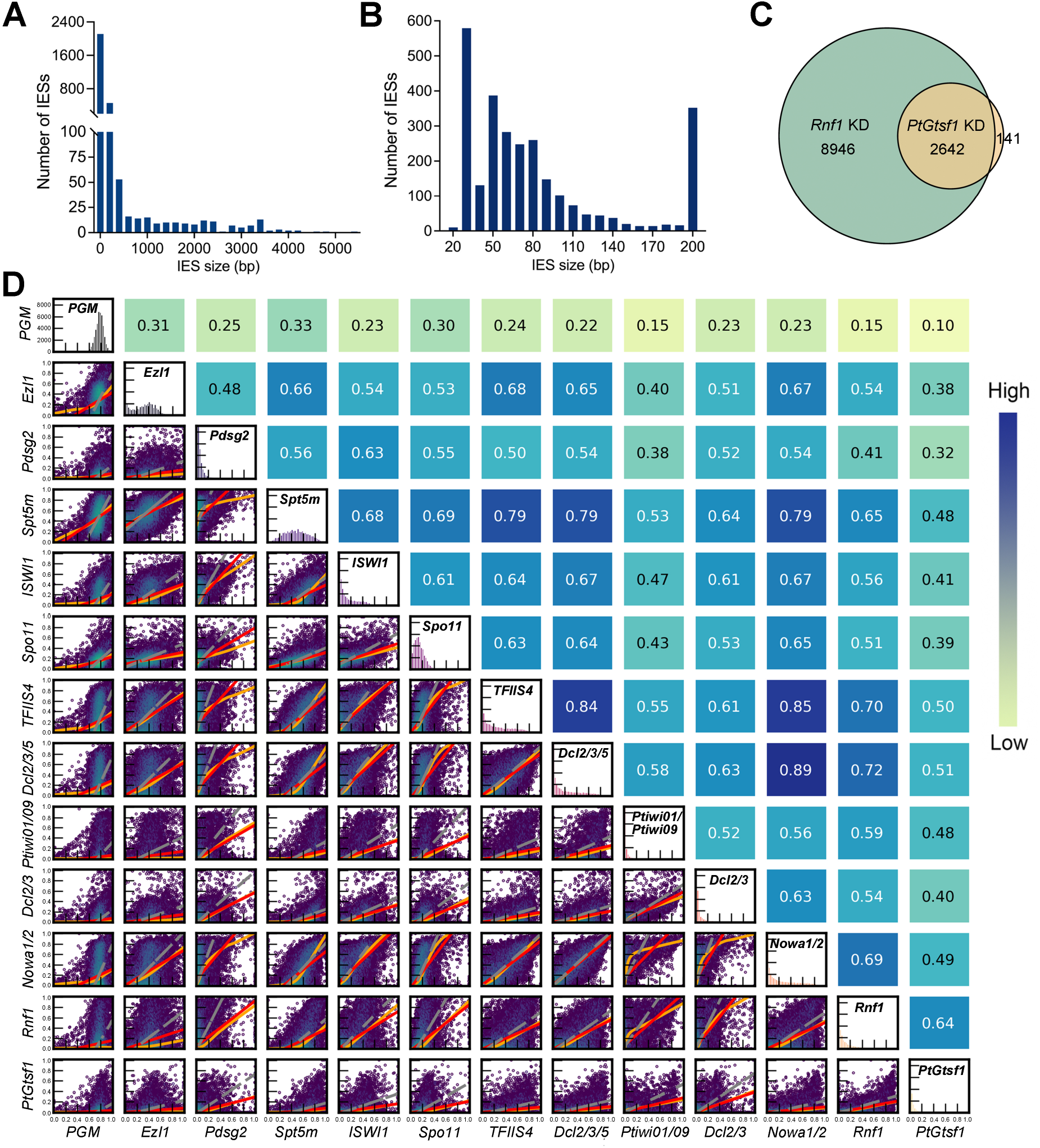
Relationship between *PtGtsf1* and other genes involved in IES elimination. A, B The size distribution of IESs (IES retention score ≥ 0.1) affected by *PtGtsf1*-KD. C Overlap of IESs retained in *PtGtsf1* and *Rnf1*-KD. D Correlation plot calculated with IES retention scores. From light green to dark blue, the correlation gradually increases.

**Figure EV3.**
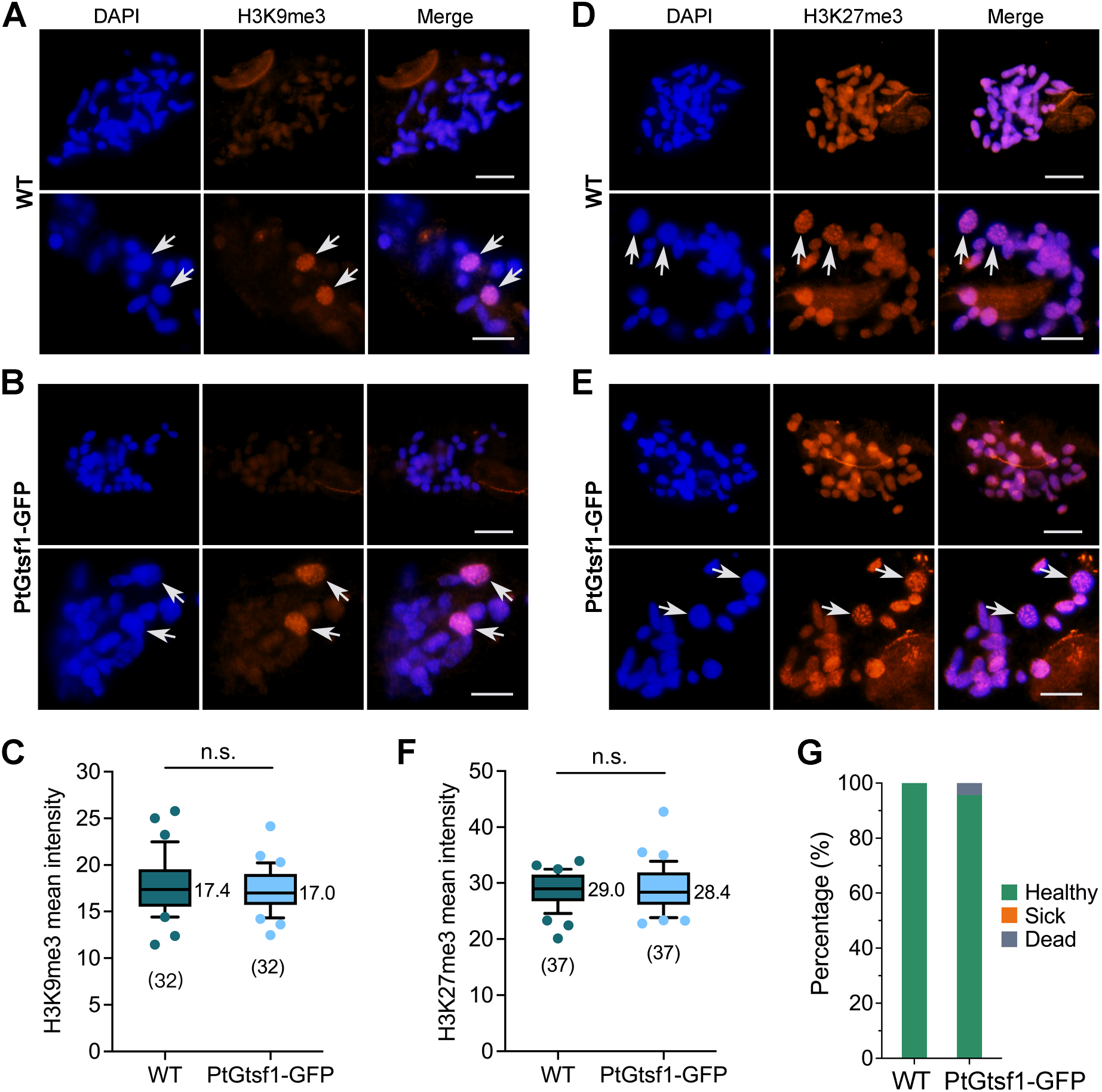
Overexpression of *PtGtsf1* does not affect the amount or distribution of H3K9me3 and H3K27me3. A, B Immunofluorescence of H3K9me3 in wild type (WT) and in cells overexpressing *PtGtsf1* (PtGtsf1-GFP). Arrows indicate new MACs. Scale bar: 10μm. C, F Mean intensity of H3K9me3 and H3K27me3 in the old MAC of WT and PtGtsf1-GFP cells. Numbers in brackets under the boxes are the individual numbers of cells used to count the signal intensity. The number to the right of the box is the median. n.s.: not significant (unpaired t-test). D, E Immunofluorescence of H3K27me3 in WT and PtGtsf1-GFP cells. G Survival test of WT and PtGtsf1-GFP cells. Cells were monitored for three consecutive days and the results of the last day are shown. Green: healthy; Orange: sick; Gray: dead.

**Figure EV4.**
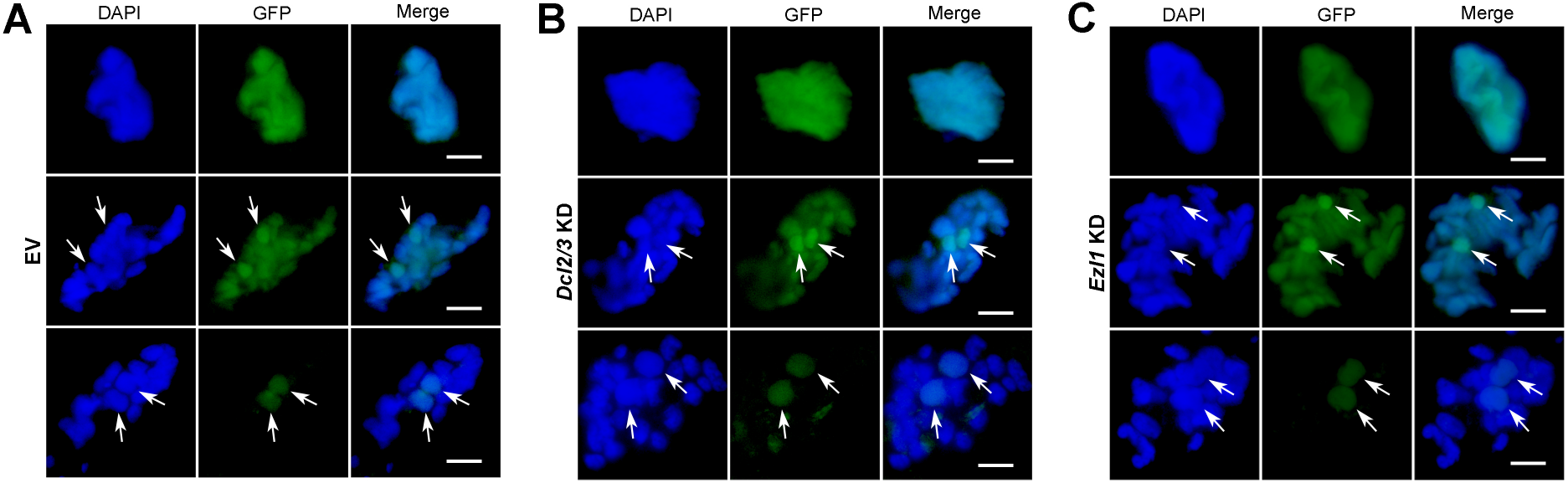
The localization of PtGTSF1 is independent of scnRNAs and PRC2. A Localization of PtGTSF1 in EV (empty vector) control. Arrows indicate new MACs. Scale bar: 10μm. B Localization of PtGTSF1 in *Dcl2/3*-KD. **C.** Localization of PtGTSF1 in *Ezl1*-KD.

**Figure EV5.**
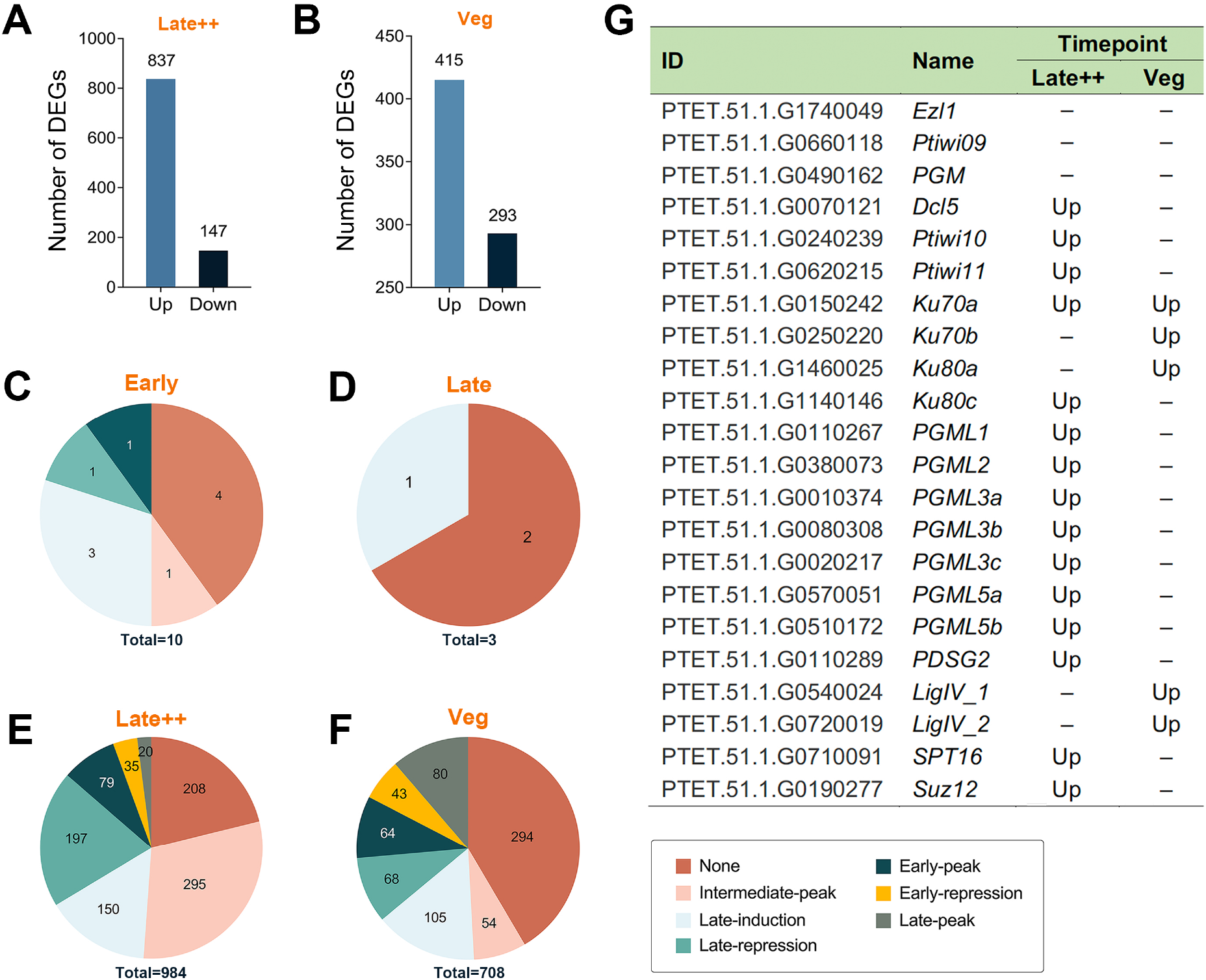
Differentially expressed genes (DEGs) in *PtGtsf1* KD. A, B Number of up and down regulated genes in *PtGtsf1* KD in Late++ and Veg stages. Up: upregulated; “–”: not differentially expressed. C-F Classification of DEGs according to their expression changes during sexual development. G Impact of *PtGtsf1* KD on the expression of genes related to genome rearrangements.

